# Guard-cell phytosterol homeostasis is critical for proper stomatal development

**DOI:** 10.1101/2024.08.29.610202

**Authors:** Chih-Chung Yen, Ya-Wen Hsu, Kuan-Chieh Leu, Sheau-Shyang Chen, Tzu-Yun Chen, Chien-Ta Juan, Chi Kuan, Jei-Fu Shaw, Chin-Min Kimmy Ho, Guang-Yuh Jauh

**Author notes:** Correspondence: Guang-Yuh Jauh and Chin-Min Kimmy Ho, Address correspondence to Guang-Yuh Jauh and Chin-Min Kimmy Ho. The author responsible for the distribution of materials integral to the findings presented in this article in accordance with the policy described in the Instructions for Authors is Guang-Yuh Jauh.

## Abstract

Stomata regulate gas exchange and control water loss in response to the environmental stimuli and their distribution in the leaf epidermis is tightly regulated during development to ensure proper patterns. Although many studies have focused on the function of early stomatal lineage cells, little is known about the role of mature guard cells (GCs) in maintaining stomatal distribution. Here, we identified a previously uncharacterized enzyme, GDSL-type sterol esterase (GSEase), that is specifically expressed in mature guard cells and catalyzes lipid droplet-stored phytosterol ester degradation. Loss of *GSEase* decreased the level of free campesterol, a biosynthetic precursor of brassinosteroids (BRs), reduced BR level, and increased stomatal density in leaves, which could be further rescued by increasing the BR signaling. Furthermore, selectively reducing the BR response in GCs by utilizing the GSEase promoter-driven *det2-1*, a mutation causing BR biosynthesis deficiency, resulted in an elevated stomatal count, as demonstrated in *gsease* plants. These results indicate that GSEase plays a critical role in maintaining phytosterol homeostasis in GCs and the released phytosterols suppress the initiation of stomatal development in adjacent cells though the BR pathway.

## Main

Stomata are microscopic valves on the epidermis that control gas exchange and water loss between the plant body and the environment ^1^. To optimize photosynthesis in a leaf, the pattern and density of stomata are well regulated during development. Stomatal precursor cells secrete epidermal patterning factor 1 and 2 (EPF1/2) to neighboring cells to prevent stomatal cell fate and regulate stomatal patterning ^2, 3, 4^. While many studies have focused on early stomatal lineage cells, the role of mature guard cells (GCs) in regulating stomatal formation remains largely elusive, even though the newly emerging stomata often appear adjacent to mature ones. Genetic analyses have shown the expression of long-chain fatty acid synthesis gene *HIGH CARBON DIOXIDE* (*HIC*) in mature GCs and the sterol reductase gene FACKEL (FK) in early stomatal lineage are required for correct stomatal patterning ^5, 6^. In addition, misregulation of the cuticle biosynthesis gene *MYB16* in early stomatal lineage caused stomatal clusters ^7^. These lines of evidence indicate lipid/sterol biosynthesis and mature stomata function in ensuring appropriate cell fate decisions. However, the mechanism of GC-derived signal to maintain the proper stomatal number in the leaf epidermis is largely unclear.

Sterols play a crucial role as a cellular membrane component and a precursor of steroid hormones and bile acids ^8^. Unlike animals or fungi, in which cholesterol or ergosterol is the unique end product of sterol biosynthesis, each plant species contains a complex mixture of sterols ^9^. Plant sterols (phytosterols) can be classified into the two most common carbon skeletons, including 29 carbons: β-sitosterol and stigmasterol, and 28 carbons: campesterol (**Sfig. 1A**) ^9^. For example, in leaves of the *Arabidopsis thaliana* ecotype Columbia, the top three abundant phytosterols are β-sitosterol (24-ethylcholesterol), stigmasterol (Δ22, 24α ethylcholesterol), and campesterol (24-methylcholesterol), for 64%, 11% and 6% proportion, respectively^10^. Phytosterols can influence cell membrane stability ^11, 12^ and are required for cell membrane plant growth, cell expansion, and division ^13^. Each phytosterol might have a unique function; for instance, β-sitosterol and stigmasterol have a major role in maintaining cell membrane structure, and campesterol acts as the precursor of brassinosteroids (BRs) ^14, 15^ ^16^.

**Fig. 1.**
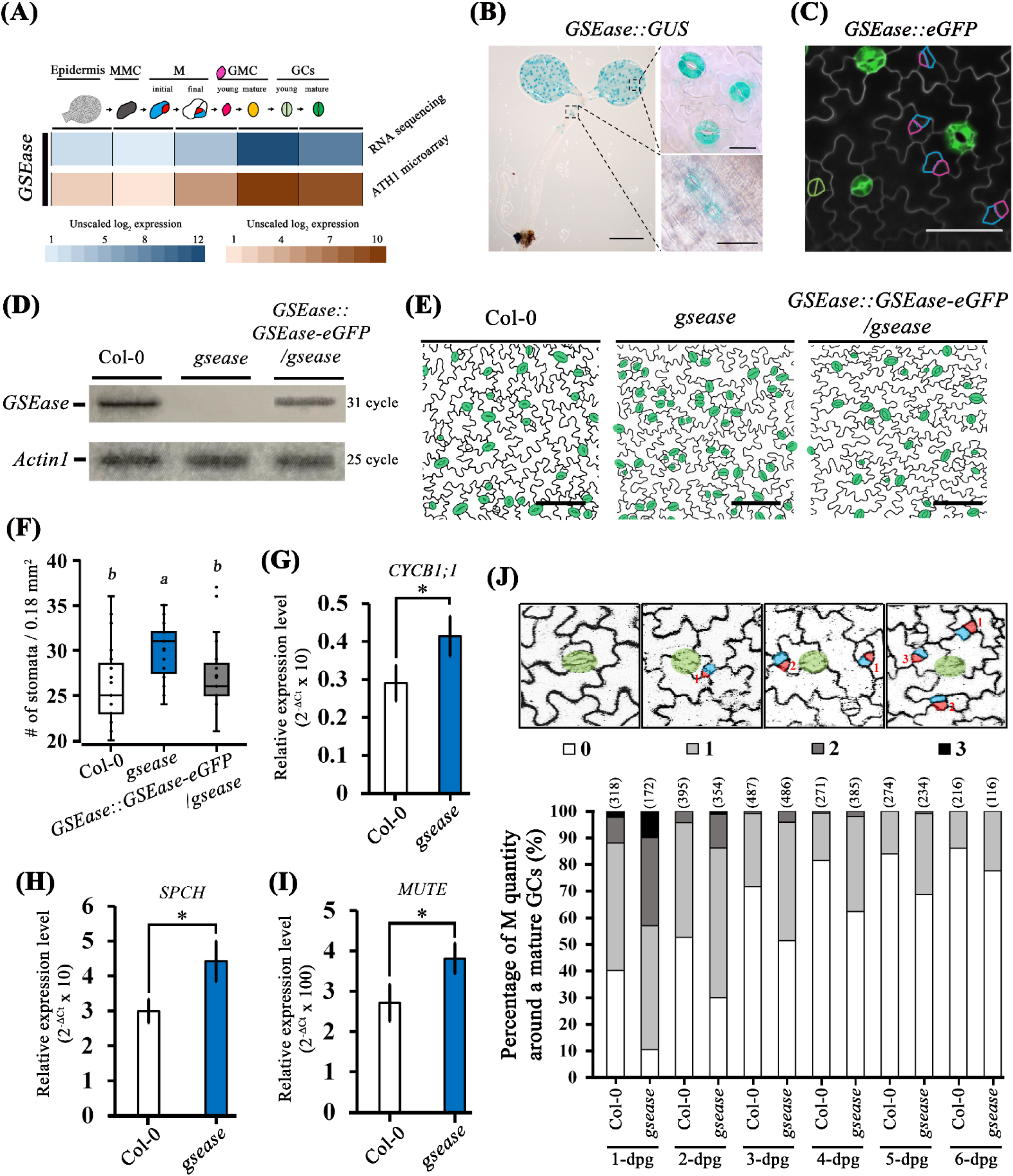
Lack of *GSEase*, a gene expressed specifically in guard cells (GCs), increases stomatal density. **(A)** Diagram showing stomata development. Epidermis, grey; meristemoid mother cells (MMCs), dark grey; initial meristemoids (Ms), red; stomatal-lineage ground cells (SLGCs), blue; young guard mother cells (GMCs), magenta; mature GMCs, orange; young guard cells (GCs, light green) and mature GCs (green). *GSEase* expression in the stomata lineage from public datasets in Adrian et al. ^31^. **(B)** Expression of *GDSL-type sterol esterase* (*GSEase*) promoter-driven *GUS* (*GSEase::GUS*) in transgenic Arabidopsis cotyledon at 5 days post-growth (dpg). GUS signal was detected in GCs of seedings (left), cotyledons (upper right), and stem (lower right). **(C)** Confocal image of *GSEase::eGFP* expression is restricted to mature GCs but not other stomatal-lineage cells. The fake outlines stained by FM 4-64FX of Ms, SLGCs and young GCs are labeled by magenta, blue and light green, respectively. **(D)** RT-PCR analysis of *GSEase* in Col-0, *gsease* and complementation line (*GSEase::GSEase-eGFP*/*gsease*). **(E)** Abaxial leaf epidermal images of Col-0, *gsease* and GSEase::GSEase-eGFP/*gsease* in 5-dpg cotyledons. Cell outline was stained with propanal iodine. GCs are labelled in green. **(F)** Quantification of stomatal density in 5-dpg cotyledons of Col-0, *gsease* and GSEase::GSEase-eGFP/*gsease*. The results were obtained from 27 independent plants (n=27), each independent plant involving one picture. Data are median (horizontal line), range (box edges) and interquartile range (whiskers). **(G-I)** qRT-PCR analysis of transcript levels of *CYCB1;1* **(G)**, *SPCH* **(H)** and *MUTE* **(I)** in Col-0 and *gsease*. **(J)** Top row shows images of 0 to 3 Ms surrounding a mature guard cell (GC). GCs, Ms and stomatal lineage cells (SLGCs) are in green, red and blue, respectively. Bottom row shows the percentage histogram of quantity of Ms around a mature GC from 1- to 6-dpg cotyledons. The number indicates total mature GCs counted. The results were obtained from more than 7 independent lines. Scale bars are 200 µm in **(B)** left, 10 µm in **(B)** right; 50 µm in **(C)** and 100 µm in **(E)**. *P*-values were calculated using one-way ANOVA with Tukey honestly significant difference test.

Phytosterol BRs negatively regulate stomatal development ^17, 18, 19^. Upon BR reception, the BR receptor BRASSINOSTEROID INSENSITIVE 1 (BRI1) dimerises with the coreceptor kinase BRI1-ASSOCIATED RECEPTOR KINASE (BAK) to phosphorylate the plasma-membrane–anchored cytoplasmic kinase BRASSINOSTEROID-SIGNALLING KINASE 1 and BRI1-SUPPESSOR1 (BSU1) phosphatase ^18^. The activated BSU1 then dephosphorylates the downstream BR-INSENSITIVE2 (BIN2) and releases its inhibition of the mitogen-activated protein kinase (MAPK) cascade, further suppressing activity of the transcription factor SPEECHLESS (SPCH), thereby inhibiting stomatal formation ^18, 20, 21, 22^. Notably, BIN2 can also regulate stomata formation via SPCH phosphorylation, independent of the MAPK pathway ^17^. Furthermore, through spatiotemporal regulation of BR signaling in asymmetrically divided stomatal precursors, BIN2 ensures the asymmetric cell fates ^23, 24^. Therefore, the function of BR signaling in early stomatal lineage cells is well studied; however, the sterol precursors of BR and their related dynamics in regulating stomatal development are unclear.

Naturally, the endogenous sterol level must be effectively regulated in a cell ^25^. Phytosterol homeostasis is a dynamic mechanism controlling the balance between phytosterol ester storage and release, preventing excess phytosterol accumulation, and satisfying the phytosterol requirement for diverse cellular events in all types of cells ^26, 27^. Free and active forms of phytosterols can be esterified at the C3-position hydroxyl group with fatty acids by acyl-CoA:cholesterol acyltransferase 1 (ACAT1) or phospholipid:sterol acyltransferase 1 (PSAT1). The resulting phytosterol esters are stored in lipid droplets (LDs) **(Sfig. 1B)** ^26, 28, 29^. The degradation of esterified sterols is catalyzed by sterol ester hydrolase, and the released free phytosterols are used for cellular metabolism. Despite playing a crucial role in maintaining the balance between the free and stored forms of sterols, the pivotal sterol ester hydrolase involved in stored phytosterol liberation has not been characterized in plants.

In this study, to identify the sterol signal regulators of stomatal development in a pre-existing stoma, we performed *in silico* screening and discovered a Gly-Asp-Ser-Leu (GDSL)-type lipase that is specifically expressed in mature GCs and whose expression is highly associated with that of genes involved in stomatal development. In an *in vitro* activity assay, the enzyme showed extremely high catalyzed activity in cholesteryl ester degradation, so we named the lipase GDSL-type sterol esterase (GSEase). In contrast to wild type plants, *gsease* mutant plants produced lower levels of “free” phytosterols with alleviated BR signaling. The *gsease* mutant consistently exhibited increased meristemoid (M) and GC density in cotyledons, and the stomata hyperproliferation phenotype could be rescued with exogenous application of BR and its precursor, campesterol. Furthermore, BR response in adjacent cell of mature GCs is decreased in *gsease*, and could be remedied by exogenous application of BR. Our results suggest that GSEase is critical in liberating phytosterol in GCs which inhibits the adjacent cells to form stomata by regulating BR biosynthesis and signaling.

## Results

### Guard-cell (GC) esterase/lipase gene *GSEase* is a negative regulator in stomatal development

Through genome-wide analysis, we previously had identified 105 GDSL-type esterase/lipase in Arabidopsis ^30^. To study the function of lipid lipase, we performed *in silico* screening to determine the tissue-specific expression patterns and possible function of the GDSL-type esterase/lipase genes. Among 105 lipases screened in eFP Browser, we found one gene*, GDSL-type sterol esterase* (*GSEase*), encoded by *At1g33811* that was highly expressed in GCs (**Sfig. 2A**), and its expression was consistent with stomata cell-type specific transcriptomes (**Fig. 1A**) ^31^. Coexpression analysis in ATTED II ^32^ (http://atted.jp) revealed 18 stomatal genes coexpressed with *GSEase,* including *STOMATAL DENSITY AND DISTRIBUTION1* (*SDD1*), *TOO MANY MOUTHS* (*TMM*), *EPF2*, *SPEECHLESS* (*SPCH)*, *SCREAM/ICE1* (*SCRM*); *SCREAM2* (*SCRM2*); *FAMA* (*FMA*); and *BREAKING OF ASYMMETRY IN THE STOMATAL LINEAGE* (*BASL*) (**Sfig. 2B**) ^20, 33^, implying the potential function of *GSEase* in stomatal development.

**Fig. 2.**
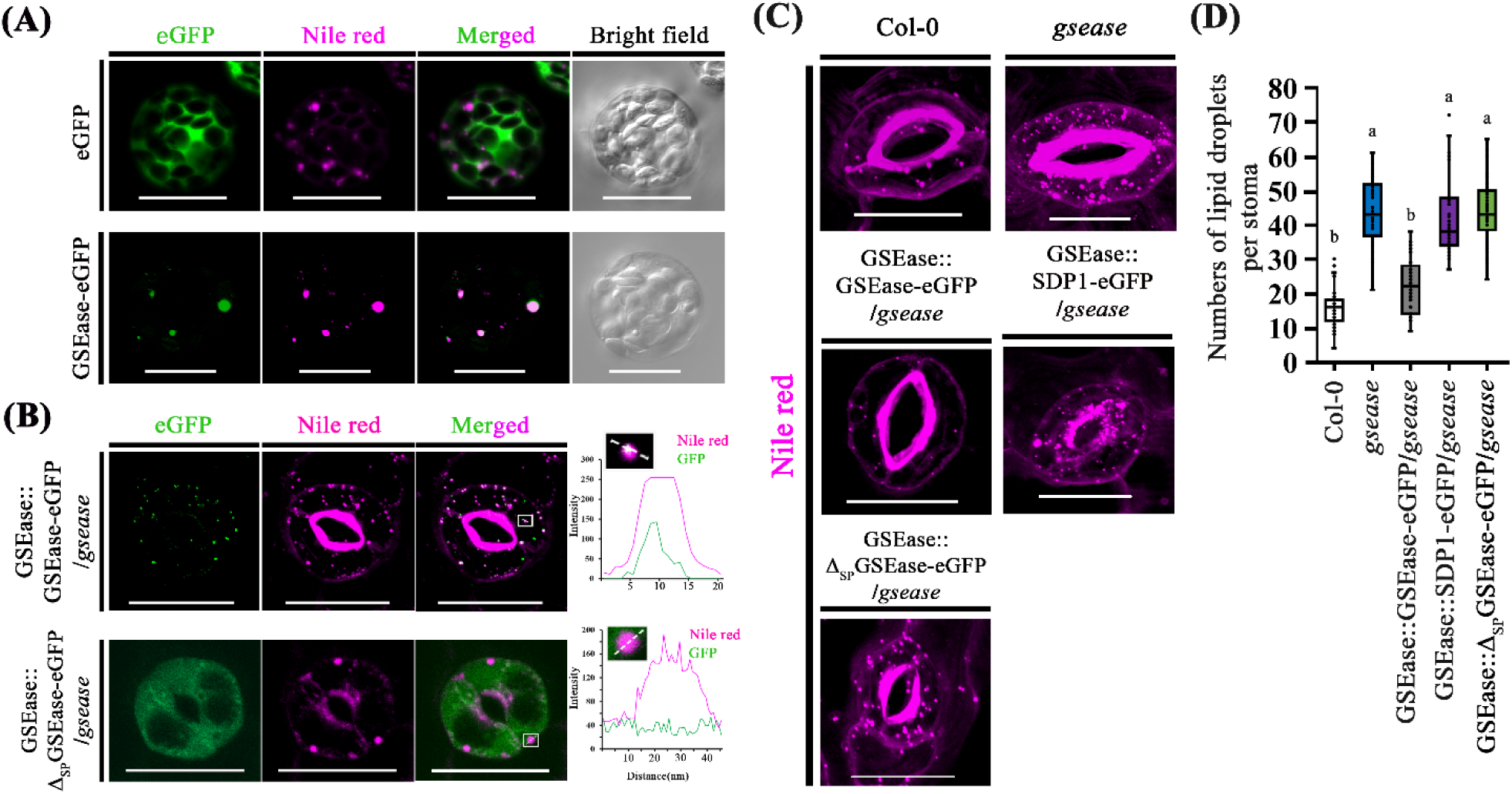
Lack of *GSEase* increased the number of lipid droplets (LDs) in stomata. **(A)** The expression of 35S promoter-driven eGFP (top) and GSEase-eGFP fusion protein (bottom) in Arabidopsis mesophyll protoplasts. **(B)** The expression of GSEase promoter-driven GSEase-eGFP fusion protein (top) and Δ_SP_GSEase-eGFP fusion protein (bottom). Right figures of fluorescent intensity, GFP signal and nile red (lipid droplets) intensity were plotted along the white dotted line in the white frame. **(C-D)** LD number in GCs in Col-0, *gsease* and its relative complementary lines. Confocal images **(C)** and quantification **(D)** of LD number of GCs in Col-0, *gsease*, GSEase::GSE-eGFP/*gsease*, GSEase::SDP1-eGFP/*gsease* and GSEase:: Δ_SP_GSE-eGFP/*gsease*. The results were obtained 45 cells (n=45) from 10 independent lines. Data are median (horizontal line), range (box edges) and interquartile range (whiskers). The signal peptide prediction is presented in **Sfig. 3**. LDs were stained with nile red and quantified. Scale bars in **(A-C)** are 20 µm. *P*-values were calculated using one-way ANOVA with Tukey honestly significant difference test.

Stomatal development begins when the multipotent meristemoid (M) mother cell (MMC) divides asymmetrically to produce a smaller M (red) and a larger stomatal lineage ground cell (SLGC, blue) ^34, 35, 36, 37^ **(Fig. 1A)**. Newly formed Ms are then converted into ellipse-shaped guard mother cells and eventually become GCs via symmetric division **(Fig. 1A)**. By contrast, SLGCs differentiate into pavement cells or generate more stomatal precursors *via* asymmetric division ^38^. As expected, in transgenic plants expressing the *GSEase* promoter-driven β-glucuronidase (GUS) (*GSEase::GUS*), GUS signals were exclusively in mature GCs in cotyledons and hypocotyls (**Fig. 1B**). Consistently, signals from green fluorescent protein (GFP) driven by the GSE promoter (*GSEase::GFP*) were also found in mature GCs but not the other stomatal lineage cells in 2-day post-germination (dpg) seedings (**Fig. 1C**).

Because GSEase was localized and predicted to function in GCs, we obtained the *gsease* T-DNA mutant, with an insertion at the fourth exon of the *At1g33811* locus (*GK-492D11*) (**Sfig. 3A**). *GSEase* gene expression was barely detected in *gsease* as compared with the wild type (Col-0) **(Fig 1D and Sfig. 3B)**.

**Fig. 3.**
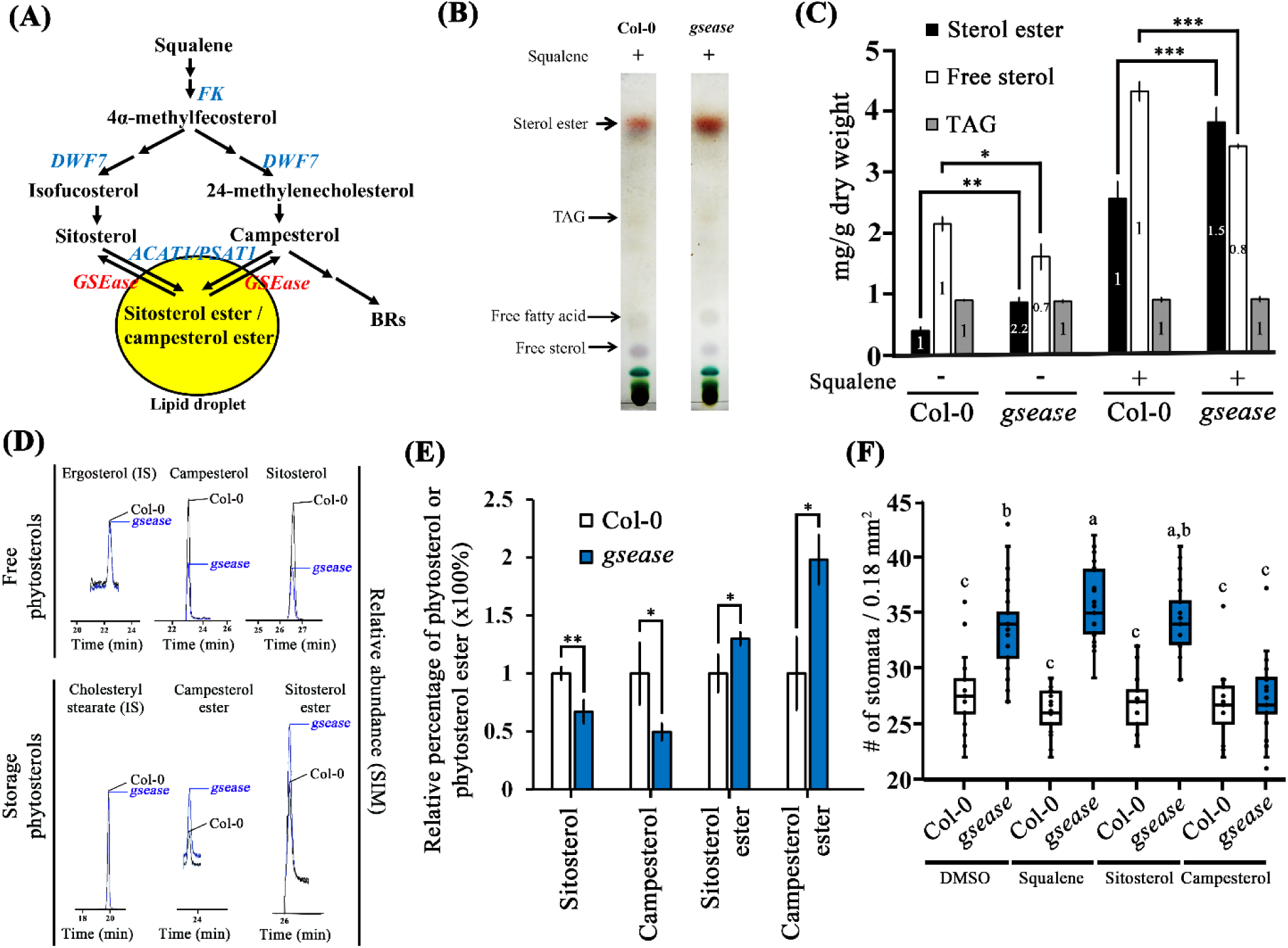
The *gsease* mutant showed reduced capability of degradation of phytosterol ester to free phytosterols. **(A)** Overview of the phytosterol homeostasis pathway. Genes related to phytosterol esters biosynthesis, *FACKEL* (*FK*); *DWARF7* (*DWF7*); *acyl-CoA:cholesterol acyltransferase 1* (*ACAT1*) and *phospholipid:sterol acyltransferase 1* (*PSAT1*), are labelled in blue. Gene related to phytosterol ester degradation, *GDSL-type sterol esterase* (*GSEase*), is labelled in red. BRs = brassinosteroids. **(B)** Thin-layer chromatography (TLC) separation of the neutral lipid fraction. Samples were collected from a rosette leaf after squalene treatment. The squalene treatment method is detailed in Methods. Neutral lipids were separated by a solvent mixture (n-hexane: diethyl ether: acetic acid, at 70:30:1, v/v) as the mobile phase. Plates were stained by immersion in a 10% sulfuric acid reagent. **(C)** Measurement of neutral lipid content in leaves of Col-0 and *gsease* plants. The number inside each bar is the ratio to Col-0, which was set to 1. Data are mean ± SD of three independent experiments. \**P*< 0.05; \*\**P*<0.01; \*\*\**P*<0.001. **(D)** Gas chromatography–mass spectrometry (GC-MS) results of free phytosterols and storage phytosterols in Col-0 and *gsease* leaves. Ergosterol (a fungi specific sterol) and cholesteryl stearate (a rarely existed sterol ester in plant ^58^) were used as the internal standard (IS) for free phytosterols and storage phytosterols, respectively. **(E)** Relative ratio of phytosterols or phytosterol esters in leaves of Col-0 and *gsease*. Data are mean ± SD of three independent experiments. \**P*< 0.05; \*\**P*<0.01. **(F)** Quantification of stomatal density with exogenous application of squalene, sitosterol and campesterol in Col-0 and *gsease*. The results were obtained from 38, 44, 22, 25, 26, 20 and 22 independent lines. Each independent plant described in **(F)** resulted in one picture. Data are median (horizontal line), range (box edges) and interquartile range (whiskers). See **Stab. 1** and **2** for quantitative results of **(C)** and **(E)**. *P*-values were calculated using one-way ANOVA with Tukey honestly significant difference test.

We further examined the stomatal phenotypes and found a significant increase in M density (**Sfig. 3CD**) and stomatal density (**Fig. 1EF**) in *gsease* mutant than wild type plants. In line with the greater stomatal phenotype in the *gsease* mutant, the transcription and translation levels of the cell cycle gene *CYCLIN B1;1* (*CYCB1;1*) were upregulated in *gsease* (**Fig. 1G**, **Sfig. 3EFG**). Levels of stomata-related genes, *SPCH* **(Fig. 1H)** and *MUTE* **(Fig. 1I)**, were also higher in *gsease* than the wild type. These results indicate elevated cell division in *gsease* plants. The stomata hyperproliferation phenotype was further complemented by the native promoter-driven *GSEase* coding sequence (CDS) **(Fig. 1DEF** and **Sfig. 4ABC)**. Hence, GSEase is a negative regulator in stomatal development.

Stomatal formation often takes place around an existing mature GC. To reveal the relation between GCs and stomata production onset, we monitored the number of Ms, an early-stage stomatal precursor cells, from 1- to 6-dpg cotyledons. About 90% Ms were found around a GC in *gsease* versus about 60% in wild type plants (89.5% vs 59.7%, respectively) when the seedings were only 1-dpg. The percentage of Ms around a mature stoma in 2- to 6-dpg cotyledons was higher in *gsease* than the wild type **(Fig. 1J)**. These observations may further indicate that the function of GSEase in stomatal development is to prevent M production around a mature GC.

### The GDSL-type sterol esterase GSEase prefers sterol ester degradation *in vitro*

Many studies have indicated that GDSL-type esterase/lipases may have flexible substrate specificity^30^. Overexpressed GSEase in an *Escherichia coli* or in a *Pichia pastoris* system was not effective in our experimental setup. Thus, to evaluate the enzyme catalytic function, we generated a *GSEase CDS* (**Sfig. 5**) overexpression line (*35S::GSEase-6x histidine*) in *A. thaliana*. GSEase recombinant protein was then purified with diethylaminoethanol (DEAE) sepharose and the histidine-tag-column procedure (**Sfig. 6A**) and verified by western blot assay (**Sfig. 6B**). The purified protein was then used for characterizing its substrate specificity *in vitro*. GSEase esterase activity was measured by using the hydrolysis of *p*-NP esters containing fatty acids of various chain lengths (C10:0 to C18:0). GSEase preferred to hydrolyze *p*-NP stearates with a long chain (C18:0) (**Tab. 1**).

**Tab 1.**
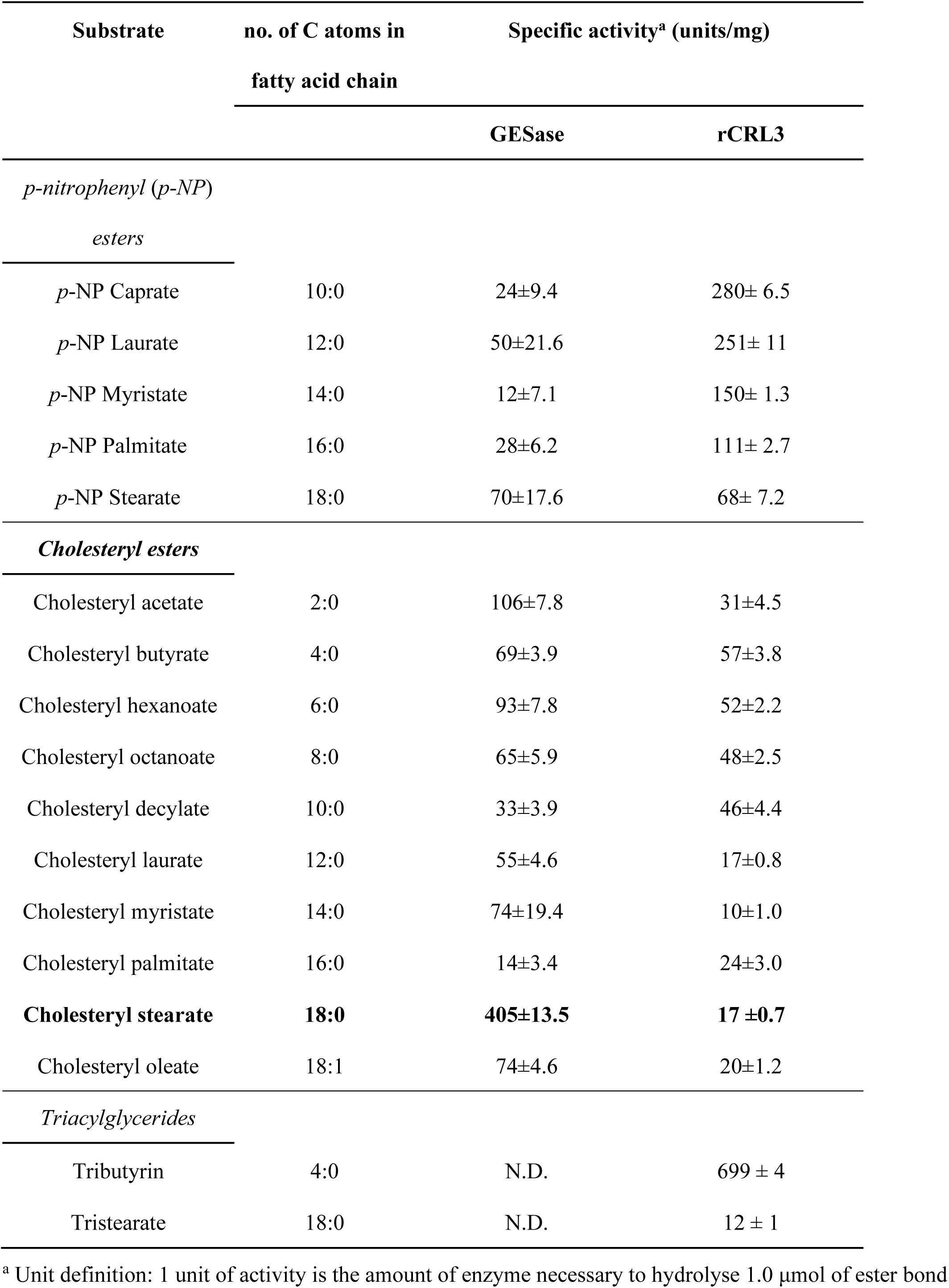
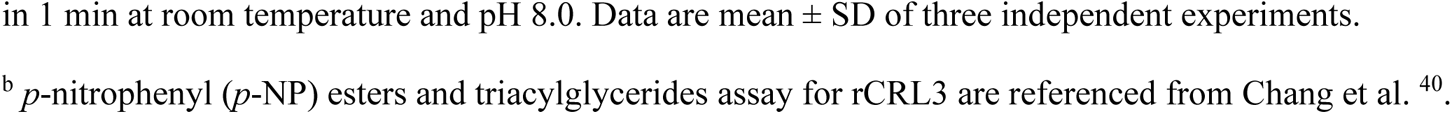
Substrate specificity of GSEase and recombinant *Candida rugosa* LIP3 lipase 3 (rCRL3) in the hydrolysis of *p*-nitrophenyl (*p*-NP) esters, cholesteryl esters, and triglycerides

| Substrate | no. of C atoms in<br>fatty acid chain | Specific activity<sup>a</sup> (units/mg) |  |
| --- | --- | --- | --- |
|  |  | GESase | rCRL3 |
| *p*-nitrophenyl (*p*-NP) |  |  |  |
| *esters* |  |  |  |
| *p*-NP Caprate | 10:0 | 24±9.4 | 280± 6.5 |
| *p*-NP Laurate | 12:0 | 50±21.6 | 251± 11 |
| *p*-NP Myristate | 14:0 | 12±7.1 | 150± 1.3 |
| *p*-NP Palmitate | 16:0 | 28±6.2 | 111± 2.7 |
| *p*-NP Stearate | 18:0 | 70±17.6 | 68± 7.2 |
| *Cholesteryl esters* |  |  |  |
| Cholesteryl acetate | 2:0 | 106±7.8 | 31±4.5 |
| Cholesteryl butyrate | 4:0 | 69±3.9 | 57±3.8 |
| Cholesteryl hexanoate | 6:0 | 93±7.8 | 52±2.2 |
| Cholesteryl octanoate | 8:0 | 65±5.9 | 48±2.5 |
| Cholesteryl decylate | 10:0 | 33±3.9 | 46±4.4 |
| Cholesteryl laurate | 12:0 | 55±4.6 | 17±0.8 |
| Cholesteryl myristate | 14:0 | 74±19.4 | 10±1.0 |
| Cholesteryl palmitate | 16:0 | 14±3.4 | 24±3.0 |
| **Cholesteryl stearate** | **18:0** | **405±13.5** | **17 ±0.7** |
| Cholesteryl oleate | 18:1 | 74±4.6 | 20±1.2 |
| *Triacylglycerides* |  |  |  |
| Tributyrin | 4:0 | N.D. | 699 ± 4 |
| Tristearate | 18:0 | N.D. | 12 ± 1 |

<sup>a</sup> Unit definition: 1 unit of activity is the amount of enzyme necessary to hydrolyse 1.0 µmol of ester bond
in 1 min at room temperature and pH 8.0. Data are mean ± SD of three independent experiments.
<sup>b</sup> *p*-nitrophenyl (*p*-NP) esters and triacylglycerides assay for rCRL3 are referenced from Chang et al.<sup>40</sup>.

Moreover, we analysed GSEase activity on the hydrolysis of cholesteryl esters of various fatty-acid chain lengths by using a peroxidase/cholesterol oxidase-coupled system ^39^. In contrast to esterase-assay findings, the cholesteryl esterase activity was significantly higher for GSEase than *Candida rugose* lipase 3 (rCRL3), a well-used enzyme for steryl ester hydrolysis in industrial applications ^40^, on cholesterol eaters (except for cholesteryl decylate and cholesteryl palmitate) (**Tab. 1**). GSEase exhibited a preference for hydrolyzing cholesteryl stearate (C18:0) (**Tab. 1**), a long-chain-length substrate, resembling the esterase-assay result for *p*-NP esters. The specific activity of GSEase on cholesteryl stearate (C18:0) (405 ± 13.5 units/mg) was 23-fold higher than that of rCRL3 (17 ± 0.7 units/mg) (**Tab. 1**). Thus, relative to rCRL3, GSEase had superior *in vitro* activity toward the hydrolysis of long-chain saturated cholesteryl esters. Notably, GSEase had a very weak preference in hydrolyzing triacylglycerols (TAGs) with tributyrin and tristearate, in contrast to rCRL3, with extremely high lipase activity. Our results suggest that GSEase is a GDSL-type sterol esterase (EC 3.1.1.13) involved in sterol ester degradation.

### *GSEase* regulates phytosterol homeostasis in lipid droplets (LDs) in GCs

Sterol esters are stored in LDs in cells ^26, 28, 29^; therefore, we predicted that the sterol esterase GSEase was located with LDs to hydrolase sterol esters and control the sterol homeostasis. To investigate the localization of GSEase, we expressed *GSEase-GFP* under a 35S promoter (35S:: *GSEase-GFP*) and its native promoter (*GSEase*:: *GSEase-GFP*) and transformed these constructs into mesophyll protoplasts (**Fig. 2A**) and *gsease* plants (**Fig. 2B**), respectively. In both cases, GSEase-GFP co-localized with nile red-stained LDs (**Fig. 2AB**). Previous studies have revealed that the deficiency of hydrolytic sterol ester enzymes or excess sterol results in larger LDs and more intracellular LDs ^41, 42, 43^. To test whether GSEase functions to degrade phytosterol esters in LDs, we compared the number of LDs in 5-dpg wild type and *gsease* seedlings. We focused on the LDs in GCs and found three times the number of LDs in *gsease* than wild type plants (**Fig. 2CD**). The phenotype of the increased number of LDs could be rescued by expressing GSEase native promoter-driven *GSEase–GFP* in *gsease* (**Fig. 2CD**). Notably, our domain analysis of GSEase (**Sfig. 5**) also indicated that the *N-terminal signal-peptide-truncated GSEase* driven by GSEase promoter (*GSEase:: _ΔSP_GSE-eGFP*) failed to target to LDs (**Fig. 2B**) and failed to rescue the *gsease* phenotype (**Fig. 2CD**). These results suggest that the N-terminal signal-peptide is required for GSEase localization and that GSEase is involved in the degradation/release of lipids in LDs.

Both phytosterol esters and TAGs are neutral lipids that are stored in LDs. The inability to degrade TAGs may also cause LD accumulation ^44, 45^ in *Arabidopsis*. To exclude the possibility that LD accumulation in *gsease* is caused by TAG accumulation, we used RT-PCR to verify the expression of two TAG-degradation related genes: *SUGAR-DEPENDENT1* (*SDP1*), a TAG lipase with TAG hydrolase activity but no cholesteryl esterase activity ^44^, and *COMPARATIVE GENE IDENTIFICATION-58* (*CGI-58*), a TAG lipase coactivator whose defect caused TAG accumulation in LDs ^45^. The expression of *SDP1* and *CGI-58* was similar in Col-0 and *gsease* leaves (**Sfig. 7A**), and the wild type and *gsease* leaves did not differ in TAG quantity (**Fig. 3C**) or fatty acid composition (**Sfig. 7B**), which indicates that LD accumulation in *gsease* is not due to altered TAG degradation.

We simultaneously tested the phytosterol ester synthesis genes *ACAT1* and *PSAT1*. *ACAT1* but not *PSAT1* was upregulated in *gsease* versus the wild type (**Sfig. 7A**), which suggests that the LD-accumulation phenotype in *gsease* may also be controlled by upregulation of *ACAT1*.

In another approach to confirm that phytosterol esters rather than TAG degradation contributed to the *gsease* phenotype, we generated a line expressing *SDP1-eGFP* driven by the GSEase promoter (*GSEase::SDP1-eGFP*) to boost TAG hydrolysis in GCs. The stomatal expression of *SDP1-eGFP* could not rescue the stomatal phenotype (**Sfig. 4ABC**) or LD number (**Fig. 2CD**) in *gsease*. These results demonstrate that the excessive LD-accumulation phenotype in *gsease* was caused by the disruption of phytosterol degradation and not TAG degradation.

### Disruption of *GSEase* significantly reduces the amount of cytosolic-free phytosterol and accumulates phytosterol esters

LDs are the major place to store sterol in its ester form. Squalene is a precursor of sitosterol and campesterol (**Fig. 3A**) in the steroid biosynthesis pathway (**Sfig. 8A**) ^43^. Instead of being free sitosterol and campesterol, they are esterified with fatty acids to sitosterol ester and campesterol ester before being stored in LDs.

Because GSEase is localized in LDs and functions as an esterase, it could be involved in phytosterol ester degradation in LDs in GCs (**Tab. 1 and Fig. 2B**). Application of squalene to *Arabidopsis* leaves increases the quantity of phytosterols, and excessive phytosterols typically are esterified to phytosterol esters before storage in LDs ^28^. Therefore, to emphasize the enzymatic functions of GSEase in plants, we treated plants with squalene to increase the quantity of phytosterol esters in LDs. First, we tested whether increased LD number and size caused by squalene treatment also occurred in GCs. After squalene treatment, LD quantity was increased in both *gsease* and wild type plants (**Sfig. 9AB**). Although the LD number was not increased in *gsease* GCs (**Sfig. 9B**), larger LDs were frequent (**Sfig. 9CD)**. The results indicate excessive phytosterols stored in the LDs of GCs.

To test whether GSEase is involved in phytosterol ester degradation, we treated plants with squalene and examined lipid composition by analyzing the total quantity of lipids extracted from 28-day-old leaves of wild type and *gsease* plants. Neutral lipids (e.g., free sterols, sterol esters, free fatty acids, and TAGs) were fractionated from polar lipids by solid phase extraction before thin-layer chromatography (TLC) separation. As compared with wild type leaves, *gsease* leaves had more sterol esters but fewer free sterols (**Fig. 3B**). The quantity of phytosterol esters was much higher in *gsease* than the wild type (0.86 ± 0.08 vs 0.39 ± 0.06 mg/g dry weight [DW]). In contrast, the amount of free phytosterols was much lower in *gsease* than wild type leaves (1.6 ± 0.20 vs 2.15 ± 0.12 mg/g DW) (**Fig. 3C** and **Stab. 1**). To exclude the possibility that the decrease in free sterols in *gsease* was caused by the altered sterol biosynthesis, we used quantitative PCR (q-PCR) to detect sterol biosynthesis genes and found no significant difference in levels of *FACKEL* (*FK*) and *DWARF7* (*DWF7*), genes involved in the biosynthesis of sterol esters, in *gsease* (**Fig. 3A**, **Sfig. 7CD**). In addition, quantification of neutral lipid composition in wild type and *gsease* plants, both treated with squalene, indicated a marked accumulation of phytosterol esters in *gsease* versus wild type leaves (3.82 ± 0.25 vs 2.56 ± 0.28 mg/g DW) (**Fig. 3C** and **Stab. 1**). By contrast, the level of free phytosterols were increased in wild type versus *gsease* leaves (4.33 ± 0.16 vs 3.43 ± 0.03 mg/g DW) (**Fig. 3C** and **Stab. 1**). These results indicate that the conversion of phytosterol esters to free phytosterol is mediated by GSEase. Taken together, the data suggest that GSEase is the key enzyme in hydrolyzing sterol esters in LDs of GCs. To determine which endogenous phytosterols were affected in *gsease*, phytosterol profiling and quantitative analysis were performed with gas chromatography–mass spectrometry (GC-MS) to quantify the phytosterol esters and free phytosterols in wild type and *gsease* plants (**Fig. 3D**); the MS spectra were presented in **Sfig. 10AB**. Notably, the levels of the two main free endogenous phytosterols, sitosterol and campesterol, were significantly decreased by 32.9% and 50.4% in *gsease* versus the wild type (1218.92 ± 182.34 and 51.8 ± 7.58 μg/g DW vs 1816.76 ± 97.7 and 104.5 ± 27.84 μg/g DW) (**Fig. 3E** and **Stab. 2**). By contrast, levels of esterified sitosterol and campesterol were increased by 30% and 98% in *gsease* versus the wild type (193.39 ± 8.46 and 17.55 ± 1.90 μg/g DW vs 148.76 ± 23.87 and 8.86 ± 2.78 μg/g DW) (**Fig. 3E** and **Stab. 2**). From these results, we conclude that disruption of *GSEase* significantly reduced the liberation of free sitosterol and campesterol from their esterified form.

### Stomata hyperproliferation phenotype in *gsease* can be compensated by an application of campesterol, a brassinosteroid (BR) precursor

To understand the relation between phytosterol homeostasis and increased stomatal density phenotype in *gsease*, we treated wild type and *gsease* leaves with squalene, sitosterol and campesterol. Squalene and sitosterol treatment were incapable of compensating the stomata hyperproliferation phenotype, but campesterol could rescue the stomatal phenotype in *gsease* (**Fig. 3F**), which suggests that the deficiency of free campsterol but not free sitosterol in *gsease* elevated stomatal division in leaves. Because campesterol is a BR precursor ^10, 46, 47^, this result also indicate that the increase in stomatal density was caused by inefficient BR production and its related BR signaling.

In general, squalene treatment resulted in a significant accumulation of both free sterol and sterol ester (**Fig. 3C**), but squalene treatment did not reduce the stomatal density in *gsease* (**Fig. 3F**). The treated squalene might be preferentially converted to sitosterol or the release of phytosterol from LDs in GCs may be important for regulating stomatal formation.

### Stomata hyperproliferation phenotype of *gsease* resulted from defective BR signaling

BR directly binds to BRI1 and activates BSU1 phosphatase via phosphorylation. Active BSU1 dephosphorylates BIN2, a negative regulator of stomatal development, releasing the inhibition of YODA (YDA) and promoting YDA-mediated suppression on SPCH. Therefore, BR signaling blocks the stomatal entry division by releasing YDA activity and phosphorylation on SPCH ^18, 21, 48^ **(Fig. 4A yellow box)**. Conversely, active BIN2 also phosphorylates BRASSINOSTEROID INSENSITIVE 1-EMS-SUPPRESSOR 1 (BES1). Phosphorylated BES1 is subjected to degradation. Therefore, BR signaling enables the promotion of BR-response genes by removing BIN2-mediated phosphorylation on BES1 **(Fig. 4A blue box)**. Campesterol is converted to the most active BR, brassinolide, by several enzymes, such as *DE-ETIOLATED2* (*DET2*), *DWARF4* (*DWF4)*, *CONSTITUTIVE AND PHOTOMORPHOGENESIS AND DWARFISM* (*CPD*), *ROTUNDIFOLIA3* (*ROT3*), and *BR-6-OXIDASE1* (*BR6ox1*) ^46, 47^ **(Fig. 4A green box and Sfig. 11A)**. Also, these BR biosynthetic genes are expressed in mature stomata according to cell-type-specific transcriptomic datasets (**SFig. 11B**) ^31^, which indicates that BR biosynthesis occurs in GCs. The expression of BR biosynthetic genes is regulated by feedback from BR signaling ^49, 50, 51^. Therefore, the aforementioned BR biosynthetic genes are upregulated under a BR-limited condition (**Fig. 4A green box**). To test whether *gsease* exhibited the BR deficiency phenotype because of a shortage of campesterol, we first monitored the expression of *DET2*, *DWF4*, *CPD*, *ROT3* and *BR6ox* in wild type and *gsease* plants by qRT-PCR. The expression of *DET2*, *DWF4*, *CPD*, *ROT3* and *BR6ox1* was significantly induced in *gsease* versus the wild type (**Fig. 4B-F**). By contrast, *SMALL AUXIN UP RNA 1 FROM ARABIDOPSIS THALIANA ECOTYPE COLUMBIA* (*Saur_AC1),* a BR-response gene ^50^, was downregulated in *gsease* versus the wild type (**Fig. 4G**). These results indicate the shortage of BR in *gsease* versus the wild type. Moreover, BES1 has both phosphorylated and unphosphorylated forms, and excessive BR application promotes the conversion of phosphorylated BES1 to its unphosphorylated active form ^52^. Thus, the phosphorylation status of BES1 can be used as a marker of BR level ^49, 50^. Higher ratios of unphosphorylated to phosphorylated BES1 represent higher BR levels in cells. In the mock treatment, this ratio was lower in *gsease* than the wild type (**Fig. 4H-I**), which demonstrated reduced BR level in *gsease*. As a positive control, wild type and *gsease* seedlings were treated with epi-BR. As expected, unphosphorylated BES1 in both the wild type and *gsease* seedlings was induced after epi-BR treatment. However, phosphorylated BES1 accumulation was greater in *gsease* leaves than wild type leaves (**Fig. 4H-I**). These results indicate that BR sensitivity was reduced in *gsease*, possibly because the mutant has a deficiency in campesterol release.

**Fig. 4.**
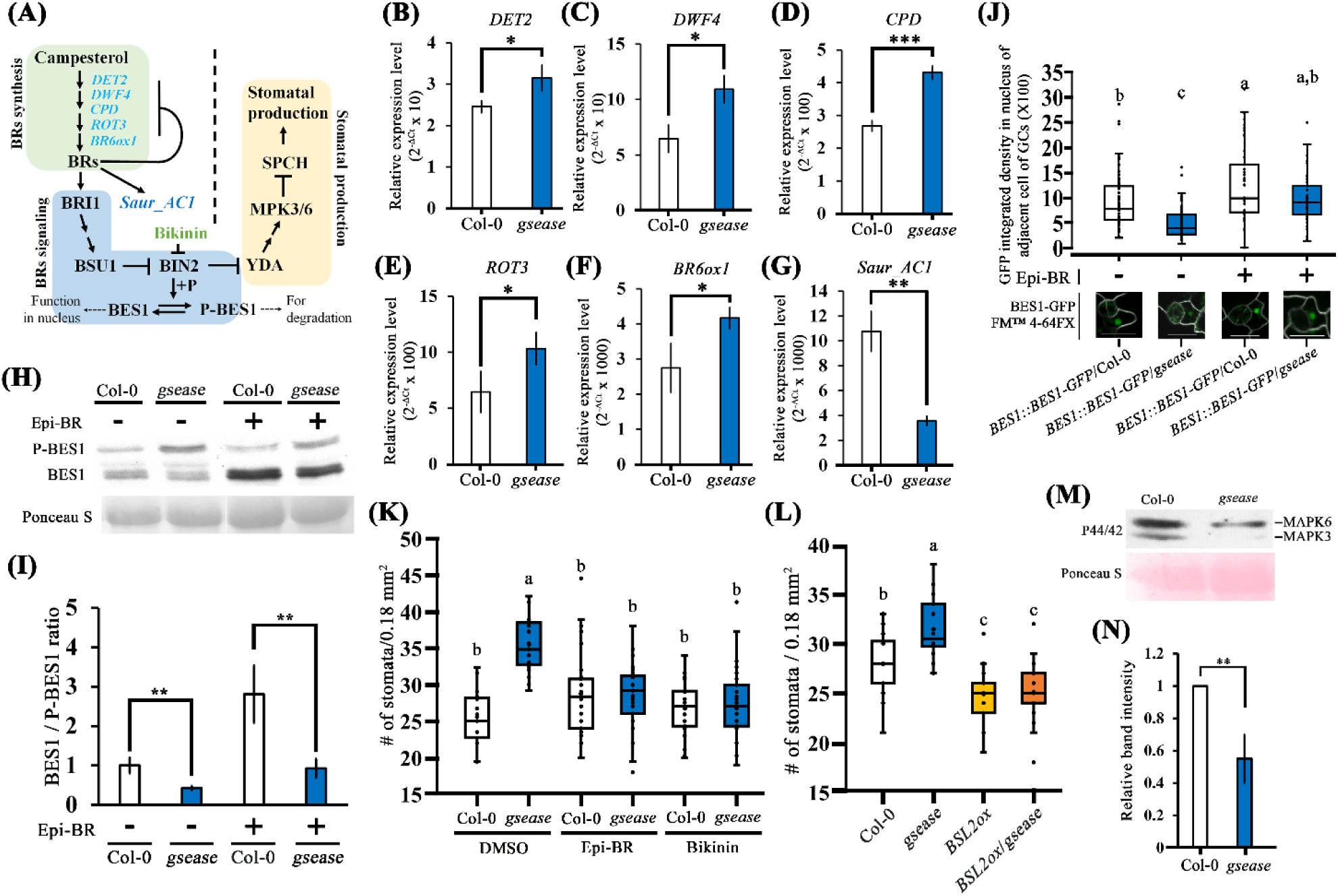
The *gsease* mutant stomata phenotype resulted from decreased BR level. **(A)** Overview of BR biosynthesis, signaling and BR mediating stomatal production pathway. Genes related to BR biosynthesis, *de-etiolated-2* (*DET2*), *DWARF4* (*DWF4*), *constitutive photomorphogenesis and dwarfism* (*CPD*), *rotundifolia 3* (*ROT3*) and *BR-6-Oxidase1* (*BR6ox1*), are labelled in light blue. BR upregulating gene, *SMALL AUXIN UP RNA 1 FROM ARABIDOPSIS THALIANA ECOTYPE COLUMBIA* (*Saur_AC1*), is labelled in dark blue. BRI1 = BRASSINOSTEROID INSENSITIVE 1; BSU1 = BRI1-SUPPESSOR1; BIN2 = GSK3-like kinase BR-INSENSITIVE2; YDA = YODA; MPK3/6 = mitogen-activated protein kinase 3/6; SPCH = SPEECHLESS. **(B-G)** Transcriptional levels of the BR biosynthesis genes and BR response gene in Col-0 and *gsease*. qRT-PCR analysis of *DET2* **(B)**, *DWF4* **(C)**, *CPD* **(D)**, *ROT3* **(E)** and *BR6ox1* **(F)** and the BR-response gene *Saur_AC1* **(G)** in 5-dpg seedlings from Col-0 and *gsease*. Data were normalized to the moderately stable reference gene *phosphoprotein phosphatase 2A subunit A3* (*PP2AA3*). Data are mean ± SD of three independent experiments. **(H-I)** Western blot analysis **(H)** and histogram **(I)** of the phosphorylation state of BRI1-EMS-SUPPRESSOR1 (BES1) in 5-dpg Col-0 and *gsease* seedlings. Protein loading was visualized with Ponceau S staining. Histogram indicates the ratio of levels of unphosphorylated BES1 (BES1) to phosphorylated BES1 (P-BES1). Western blot analysis of BES1 by using BES1 antibody (R3489-2). Data are mean ± SD of three independent experiments. \*\**P*<0.01. **(J)** Box plot is quantification of nuclear BES1pro:: BES1-eGFP fluorescence intensity in nucleus of adjacent cells of GCs in Col-0 and *gsease*. The results were obtained from 65, 72, 45, and 45 cells of more than 15 independent plants. **(K)** Quantification of stomatal density of exogenous application of epi-BR and bikinin in Col-0 and *gsease*. The results were obtained from 19, 19, 29, 26, 26 and 29 independent lines. **(J, K, L)** Data are median (horizontal line), range (box edges) and interquartile range (whiskers). **(L)** Quantification of stomatal density of overexpression of BSL2 (BSL2ox) in Col-0 and *gsease*. The results were obtained from 20, 24, 25 and 25 independent lines. Each independent plant described in **(K)** and **(L)** involved only one picture. **(M-N)** MAPK activity was decreased in *gsease* mutants. Western blot analysis **(M)** and quantification **(N)** were performed to evaluate MAPK3/6 kinase activity with an anti-phospho-ERK1/2 (pThr202/Tyr204) antibody. Protein loading was visualized with Ponceau S staining. Quantification of MAPK3/6 activity in 5-dpg Col-0 and *gsease* cotyledons (lower) involved western blot analysis. Data are mean ± SD of three independent experiments. *P*-values were calculated using one-way ANOVA with Tukey honestly significant difference test.

**Fig. 5.**
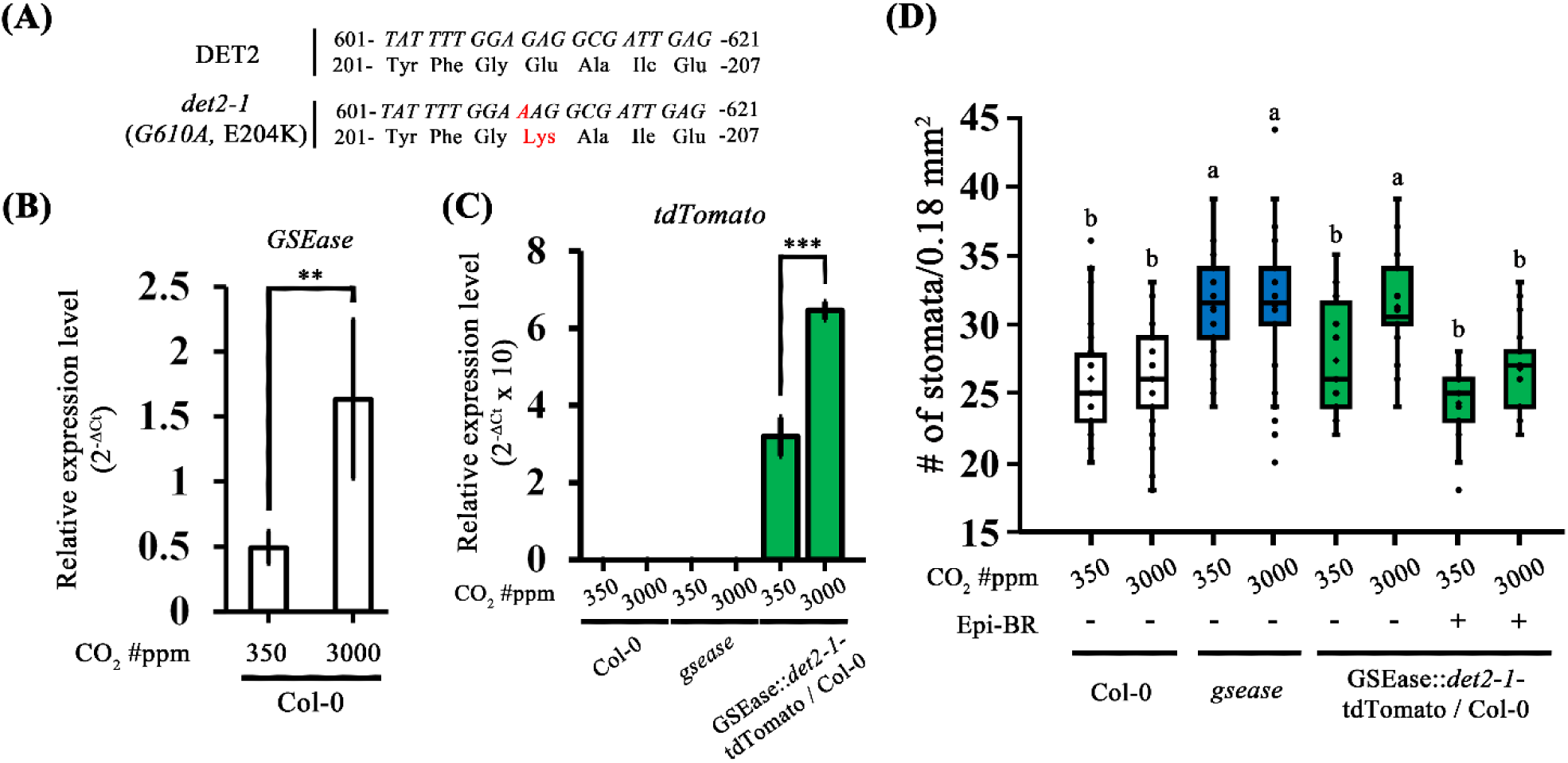
Decreasing BR level of GCs increased stomatal density. **(A)** Schematic diagram of *det2-1*, a BR biosynthesis mutation of DET2. **(B)** *GSEase* expression was elevated by high CO_2_ (3000 ppm). Data were normalized to the moderately stable reference gene *PP2AA3*. **(C)** Transcriptional levels of *det2-1-tdTomata* in Col-0, *gsease* and GSEase::*det2-1*-tdTomato / Col-0 under normal and high CO_2_ condition. **(D)** Quantification of stomatal density of Col-0, *gsease* and GSEase::*det2-1*-tdTomato / Col-0 under normal and high CO_2_ or without and with epi-BR treatment. Data are median (horizontal line), range (box edges) and interquartile range (whiskers). The mean ± SD data in **(A)** were calculated from four independent experiments and in **(C)** from three independent experiments. The results in **(D)** were obtained from 34, 22, 36, 40, 18, 22, 20, 21, 22 and 21 independent lines. Each independent plant described in **(D)** involved only one picture. *P*-values were calculated using one-way ANOVA with Tukey honestly significant difference test.

Because the activation of BR signalling results in the phosphorylation and degradation of BES1 (**Fig. 4A**), the signal of BES1-eGFP could serve as a proxy for BR level. Therefore, we monitored the effect of BR in GC adjacent cells by using the signal intensity of BES1-GFP. Transgenic plants expressing *BES1::BES1-eGFP* had similar expression for both the wild type and *gsease* (**Sfig. 12A**), but the BES1/phosphorylated-BES1 ratio was decreased in *gsease* using a GFP antibody (**Sfig. 12B**). This result illustrated that the transcription level of *BES1-eGFP* was equal in the wild type and *gsease*, but the BR level was lower in *gsease* (similar to **Fig. 4H).** Furthermore, we investigated BES1-eGFP intensity in the nuclei of cells next to GCs. The BES1-eGFP intensity in the nucleus of adjacent cells of GCs was significantly lower in *gsease* than the wild type (**Fig. 4J**). Notably, epi-BR treatment increased the BES1-eGFP intensity in GC neighboring cells in *gsease* comparable to the wild type (**Fig. 4J**), which implies that in *gsease*, the reduced BES1-eGFP intensity in cells adjacent to GCs was a result of insufficient BR. Given the function of GSEase, these data indicate that GSEase released campesterol, a BR precursor, in GCs to prevent the stomatal fate in adjacent cells.

Furthermore, to confirm that the stomatal phenotype in *gsease* was indeed due to the defect in BR signaling, we applied plants with epi-BR or bikinin (a direct BIN2 inhibitor ^18^). As expected, the stomatal phenotype in *gsease* could be rescued by both epi-BR and bikinin treatments (**Fig. 4K**). In addition, we examined the genetic interactions between *gsease* and BR signaling mutants. The stomatal phenotype in *gsease* could be eliminated by overexpressing the positive BR signaling component BSL2, a member in the BSU1 family ^53^ (**Fig. 4L**). BR regulates stomatal development by activating MAPK ^18^ (**Fig. 4A yellow box**). Thus, we also measured MAPK3/6 activity by using a p44/42 MAPK antibody in wild type and *gsease* plants and observed a decrease in MAPK3/6 activity in *gsease* (**Fig. 4M-N**). These findings illustrate that the presence of the stomatal phenotype in *gsease* is attributable to the reduced BR signalling. GSEase functions to liberate campesterol in GCs and suppresses stomatal formation in the neighbouring cells via the BR-mediating signalling pathway.

### BR formation in GCs prevents hyperproliferation of stomata

To further evaluate whether BR production in GCs is critical in regulating stomatal distribution, we constructed GSEase promoter-driven *det2-1* (G610A, E204K), a point mutation of DET2 (**Fig. 5A)** that fused *tdTomato* (*GSEase::det2-1-tdTomato*), then transferred it into the wild type plants. The *det2-1* is a BR-deficient mutant ^54^, which we designed to decrease BR level in GCs by competing with normal DET2 in the BR biosynthesis pathway **(Fig. 4A green box and Sfig. 11A)**.

In the coexpressed gene analysis, *GSEase* was highly coexpressed with genes (*ALMT12*, *HT1*, and *MPK12*) related to GC carbon dioxide (CO_2_)-response mechanisms (**Stab. 3**) ^55^, which suggests that GSEase expression might be altered by CO_2_ concentration. As predicted, *GSEase* was upregulated under elevated CO_2_ (3000 ppm) as compared with the normal condition (350 ppm) (**Fig. 5B**). However, the expression of *DET2* was not affected under the same treatment (**Sfig. 13A**). To evaluate the effect of BR signaling produced in GCs to the adjacent cells, we first expressed *det2-1-tdTomato* using a GSEase promoter. As expected, the signal of *tdTomato* was upregulated in the wild type under elevated CO_2_ (**Fig. 5C**). Then we used this line to specifically attenuate the BR production in GCs under elevated CO_2_ and monitor the stomatal phenotype. Elevated CO_2_ did not alter the stomatal phenotype in the wild type and *gsease* (**Fig. 5D**). However, increased CO_2_ concentration (3000 ppm, which activates the expression of *det2-1*), which was confirmed by elevated expression of CO_2_-treated marker *PDF1.2* (**Sfig. 13B)**, increased the stomatal density in *GSEase::det2-1-tdTomato*/Col-0, indicating the elimination of BR in GCs did influence the stomatal production. The increased stomatal density in the upregulated *det2-1-tdTomat* plants was further eliminated by epi-BR treatment under elevated CO_2_ (3000 ppm) (**Fig. 5D**). These results indicate that the BR level in GCs plays a role in controlling stomatal formation in Arabidopsis.

## Discussion

Sterol homeostasis is a critical mechanism that regulates the balanced synthesis and degradation of sterol esters in all species. Sterol ester hydrolase, an enzyme for sterol ester degradation, liberates sterols as building blocks for membrane formation and also provides cholesterol for hormone synthesis in mammalian adrenal cells and Leydig tumor cells ^26^. In humans, defective sterol ester hydrolase, such as lysosomal acid cholesterol ester hydrolase, is related to the Wolman and cholesterol ester storage diseases^26^. In yeast, sterol ester hydrolase triple-mutant *yeh1Δyeh2Δtgl1Δ* lost its viability during the stationary phase ^42^. These reports suggest that sterol esterase is involved in sterol hormone homeostasis and fundamental cell growth in mammals and yeast.

There are 105 members in the GDSL-type esterase/lipase family ^30^, and 15 members (including *GSEase*) highly expressed in GCs ^56^. The mutant phenotypic identification indicated GSEase and its 3 of homologous played redundant roles in stomatal density, morphology, and dynamics ^56^. Additionally, *GSEase* expression was up-regulated when plant grow under elevated CO_2_ condition (**Fig. 5B**), that indicates GSEase is capable to respond environment adaptation. There results implied that phytosterol homeostasis might play a role in stomatal patterning and stomatal dynamics to adapt to the environment changes, which will also need to be examined in detail. Therefore, we plan to contribute efforts to tackle the studies of coordination in phytosterol homeostasis and environment adaption in the next phase.

In this study, we found GSEase possessed activity for sterol ester degradation *in planta*. *In vitro*, GSEse exhibited extremely high substrate specificity and high activity of cholesteryl ester degradation (even more than the well-known cholesterol esterase rCRL3), thus illustrating its enzyme engineering and industrial potential. *In vivo*, GSEase releases phytosterols from LDs, and the storage and liberation of a BR biosynthesis precursor, campesterol, from LDs suggests that the storage and liberation of campesterol are potentially involved in regulating endogenous BR levels in Arabidopsis. BR homeostasis seems also mediated by phytosterol homeostasis. BR metabolism, such as esterification, is an important mechanism in the regulation of BR homeostasis. Certain BR esterified forms are present in some plant species ^50, 57^, but the storage location of esterified BR is unknown and which enzymes are involved in the liberation of esterified BR is unclear. Because the chemical structure of esterified BR is similar to that of esterified phytosterol, LDs may take esterified BR as well. Therefore, we do not exclude that GSEase is involved in the degradation and liberation of esterified BR. Unfortunately, we failed to detect the endogenous BRs in Arabidopsis seeding by ultra-performance liquid chromatography mass spectrometry due to matrix effects, let alone BR derivatives. However, we are still only beginning to understand the regulation of phytosterol homeostasis and its biological role in cells.

Treatment with squalene elevated plant-leaf sterol levels in both free and storage form (**Fig. 3C**). Notably, the increase in free phytosterol level, which accumulated because of squalene treatment, could not rescue the *gsease* phenotype. However, exogenous campesterol treatment could suppress the increased stomatal density in *gsease*, which suggests that the *gsease* stomata hyperproliferative phenotype is mainly produced in the absence of campesterol. We hypothesized that 1) squalene treatment promotes the production of other phytosterols, such as sitosterol but not campesterol and 2) all excess phytosterol from squalene is stored in LDs first, where phytosterol liberation is key to maintaining the balance between the free and ester forms of phytosterol. Details of the regulation of phytosterol homeostasis remain for further studies.

Our results provide a new model of GSEase playing a role in hydrolyzing sterol esters and maintaining phytosterol homeostasis in mature GCs. The phytosterols released from LDs may be converted to BRs and sent out from GCs. BRs are considered to be transported from the cell where they are generated to be perceived at the surface of adjacent target cells ^58^. The implication is that plant cells may influence adjacent target cells by adjusting BR production. Even recent research indicated that BRs are transported through plasmodesmata ^59^, but mature GCs are considered to lack plasmodesmata ^60^; thus, how BRs of GCs affect adjacent cells remains unclear. Secreted BRs may be received by their receptors on the surface of adjacent cells, providing a signal to activate BR-mediated MAPK signaling to suppress stomata formation in adjacent cells into a stomatal lineage (**SFig. 14**). With loss of *GSEase*, phytosterol esters (especially campesterol ester) accumulate in LDs, thus decreasing the level of free phytosterols in GCs. Decreased BRs lead to stable BIN2 to suppress MAPK signaling to increase M formation in adjacent cells (**SFig. 14**). However, we cannot rule out that phytosterols directly regulate stomata development in adjacent cells via an unknown mechanism or whether BR is synthesized in GCs or adjacent cells after phytosterols are transported. Although our findings reveal the potential effect of phytosterol on BR homeostasis, the detailed mechanism underlying phytosterol in BR biosynthesis in GCs and their corresponding cell-to-cell movement requires further investigation.

## Methods

### Plant materials and growth conditions

*Arabidopsis thaliana* (Columbia) seeds were surface-sterilised with chlorine gas for 3 h. After stratification at 4°C for 2−3 days, seeds were plated in ½ Murashige-Skoog medium (pH 5.7) containing 1% sucrose and 0.8% phyto agar, then grown inside a growth chamber on a 16-h light/8-h dark cycle at 22°C. The *gsease* mutant (GK-492D11-019647) was obtained from the *Arabidopsis* Biological Resource Center (Ohio State University, Columbus, OH, USA). The *gsease* mutant was verified as homozygous for the T-DNA insert in the GSEase locus by PCR genotyping, which was performed according to the method outlined in the GABI-Kat database (http://www.gabi-kat.de/db/showseq.php?line=492D11). High CO_2_ condition was set growth chamber on a 16-h light/8-h dark cycle at 22°C with 4000ppm CO_2_. Primers are in **Stab. 4**.

### DNA manipulation

The GSEase promoter region (1262 bp) was amplified from plant genomic DNA. The GSEase promoter region fused to the *GUS* gene was produced in pBI121. *GSEase::eGFP* was produced by replacing GUS with enhanced green fluorescent protein (eGFP). The *At1g33811.1* (*GSEase*) open reading frame was amplified by PCR. Translational fusions of a 35S promoter driving *GSEase-6x histidine* were synthesised in pBI121. The GSEase promoter driving *GSEase–eGFP* (*GSEase::GSEase-eGFP*) was synthesised by inserting the *GSEase* CDS into *GSEase::eGFP*. *GSEase::SUGAR-DEPENDENT1 (SDP1)-eGFP* and *GSEase:: Δ_SP_GSEase-eGFP* were constructed by replacing *GSEase* cDNA with *SDP1* CDS and signalling peptide–truncated *GSEase-eGFP* (*Δ_SP_GSEase*) cDNA, respectively. *CYCB1;1:: 3NLS-mCherry* and *BES1::BES1-eGFP* were produced in pPZP221. The primer design depended on a sequence from The Arabidopsis Information Resource (TAIR). All primers used in this experiment are in **Stab. 4**.

### Plant transformations

All constructs were introduced into wild type or *gsease* leaves of *A. thaliana* via floral dipping. *Agrobacterium tumefaciens* strain GB3101 was used for transformation. A 50-mL overnight *A. tumefaciens* culture in a lysogeny broth medium was centrifuged at 6000 *g* for 10 min at room temperature before being resuspended in a 50-mL sucrose solution (5% sucrose, 0.005% Silwet L-77). Flowers from 5- to 7-week-old plants grown in soil under long-day conditions (16-h light, 8-h dark) were dipped in the *A. tumefaciens* solution for 30 s and left to dry. The procedure was performed three consecutive times at 1-week intervals.

### GUS staining

Whole seedling plants were immersed in a GUS staining solution (containing 100 mM NaH_2_PO_4_ [pH 7.0), 10 mM EDTA, 0.5 mM Triton X-100, 0.5 mM K_4_[Fe(CN)_6_], and 2 mM 5-bromo-4-chloro-3-indolyl-β-glucuronide), then incubated for 4 h at 37°C. The staining solution was removed and samples were washed with 70% ethanol.

### Quantification of stomatal phenotype and protein sub-cellular localization visualization

For stomatal phenotype assessment, seedlings at 1 to 6 days post-growth (dpg) were stained with propidium iodide, FM 4-64FX and nile red. To visualize GSEase-eGFP, the protoplasts and seedings were treated with nile red for 10 min. Images were acquired under a laser scanning confocal microscope (Zeiss LSM 510 Meta Confocal Microscope, Carl Zeiss, Germany) and analysed with a Zeiss LSM imaging browser. Similar central regions in the epidermis were captured to minimise variation. Propidium iodide and FM 4-64FX were acquired from Invitrogen (Waltham, MA, USA); nile red was acquired from Sigma (St. Louis, MO, USA).

### Imaging LDs

Cotyledons were stained with 6.5 mg/mL nile red. LD number and area were quantified by using the LSM Image Browser Z-axis stack and ImageJ software (National Institutes of Health, v1.43T).

## Protein purification

Two-week-old homozygous seedlings were harvested to extract GSEase protein for purification. GSEase-6x histidine were overexpressed by the 35S promoter. The plants were ground in liquid nitrogen, and the product was resuspended in lysis buffer containing 50 mM Tris-HCl (pH 8.0), 300 mM NaCl, and an EDTA-free protease inhibitor cocktail. The homogenates were spun at 4000 *g* for 30 min, and the supernatant was recentrifuged at 22,000 *g* for 90 min. The supernatant was filtered through a 0.45 μM Millipore membrane. Subsequently, the crude cell lysate (containing overexpressed recombinant GSEase protein) was loaded onto a DEAE Sepharose Fast Flow column and HisTrap Ni Sepharose column (GE Healthcare, Inc., Chicago, IL, USA). All protein manipulations were performed at 4°C. The column was washed with different-concentration imidazole to elute the protein. Fractions were collected and analysed on 12% SDS-PAGE gel and stained with Coomassie Blue staining. The GSEase with 6X His tag was approximately 41 kDa, similar to the predicted mature protein molecular weight on the compute pI/Mw tool website (https://web.expasy.org/compute_pi/). A monoclonal 6X His tag antibody (GTX359, GeneTex, USA) was used for detecting the GSEase-6X His in a western blot assay. The expression and purification of *Candida rugusa* lipase 3 (CRL3) were as reported ^40^.

### Enzyme substrate specificity assay

For the esterase activity assay, the substrate solution was 2.5 mM of *p*-nitrophenyl (*p*-NP) esters: *p*-NP caprate (C10:0), *p*-NP laurate (C12:0), *p*-NP myristate (C14:0), *p*-NP palmitate (C16:0), and *p*-NP stearate (C18:0) (Sigma, St. Louis, MO, USA) dissolved in a 50-mM sodium phosphate buffer with pH 8.0, containing 2.1% (v/v) Triton X-100. The enzyme reactions were measured at room temperature, and a 10-min reaction was initiated by adding a 2.0-μL enzyme solution into a 200-μL substrate solution. Enzyme activity was determined according to absorbance changes at 405 nm. The molar extinction coefficient of *p*-NP in this buffer system was 1725 M^−1^ cm^−1^. For the cholesterol esterase activity assay, the cholesterol esterase activity of GSEase was analysed with a peroxidase/cholesterol oxidase coupled system to measure the formation of cholesterol and fatty acid during the hydrolysis of cholesteryl esters of various chain lengths: cholesteryl acetate (C2:0), cholesteryl butyrate (C4:0), cholesteryl hexanoate (C6:0), cholesteryl octanoate (C10:0), cholesteryl n-decylate (C10:0), cholesteryl n-laurate (C12:0), cholesteryl n-myristate (C14:0), cholesteryl n-palmitate (C16:0), cholesteryl n-stearate (C18:0), and cholesteryl n-oleate (C18:1) (Sigma, St. Louis, MO, USA). The hydrolytic reaction was conducted at room temperature for 1 h in a mixture of 200 μL reagent solution: 100 mM Tris/HCl (pH 8.0), 50 mM MgCl_2_, 6 mM phenol, 1 mM 4-aminoantipyrine, 4 mM 3,4-dichlorophenol, 10 mM sodium cholate, 3 g/L Thesit (Boehringer Mannheim, Mannheim, Germany), 500 units/L cholesterol oxidase (Sigma, St. Louis, MO, USA), 400 units/L peroxidase (Sigma, St. Louis, MO, USA), and 100 μL of 100 g/L Thesit solution containing 10 mM substrate. The absorbance was recorded at 500 nm. One unit of cholesterol esterase activity was defined as the quantity of enzyme that hydrolyses 1.0 μmoL cholesteryl esters to cholesterol and fatty acids in 1 min. Triacylglycerol (TAG) activity was as reported ^61^.

### Enzyme kinetic assay

The kinetic studies were performed at room temperature, and enzyme activities were measured spectrophotometrically. For the esterase activity assay, six concentrations of *p*-nitrophenyl stearate (0.15625, 0.3125, 0.625, 1.25, 2.5, 5 mM) and cholesteryl stearate (0.625, 1.25, 2.5, 5, 10 and 20 mM) were prepared by dissolving *p*-NP butyrate and cholesterol esterase buffer.

### Squalene treatments in soil

Squalene treatments involved 13-day-old plants; 10 µL squalene was irrigated thrice at 5-day intervals. Rosette leaves were collected at approximately 2 days after the third application and frozen. The collected samples were used for the following step. Squalene was acquired from Sigma (St. Louis, MO, USA).

### Neutral lipid extraction

Plant materials were obtained from 28-day-old plants and frozen in liquid nitrogen. Total lipids were extracted from plant tissues using the chloroform/methanol method. Each sample up to 0.1 g dry weight was covered with 3 mL preheated isopropanol at 75°C for 15 min before being cooled to room temperature. We then added 1.5 mL chloroform and 0.6 mL water, then incubated with shaking for 1 h. After transferring the lipid extract to a fresh tube, we extracted tissue twice with 4 mL of chloroform:methanol (2:1 v/v). To combine lipid extracts, we added 1 mL of 1 M KCl to separate the two phases before discarding the upper phase (polar phase). Subsequently, we added 2 mL water, then vortexed, centrifuged, and discarded the upper phase (polar phase). The nonpolar phase was dried under a stream of nitrogen. Nonpolar samples were dissolved in 1 mL chloroform:MeOH (95:5, v/v). Neutral lipids were further separated from polar lipids by silica gel column chromatography (Supelco Discovery DSC-Si 6 mL, 500-mg solid phase extraction cartridges) in chloroform:MeOH (95:5, v/v).

### Thin-layer chromatography (TLC) and lipid analysis

Neutral lipids were analysed by TLC. Samples were separated by using BAKER Si250-PA(19C)-Silica gel (J.T. Baker) and a solvent mixture of n-hexane: diethyl ether: acetic acid (70:30: 1, v/v) with cholesteryl oleate, trioleylglycerol, oleic acid, and cholesterol as references. Plates were stained by immersion in a 10% sulfuric acid reagent or 0.01% primuline, then heated and viewed under an ultraviolet lamp.

### Quantitative analysis of neutral lipids

Individual samples were scraped from the TLC plate. Then we extracted them in 200 µL chloroform:methanol (2:1 v/v) before drying under a stream of nitrogen. Sterol esters were hydrolysed by heating under reflux for 1 h with methanolic KOH (6%). The sterols were then extracted three times with hexane. Both free sterols and sterols corresponding to sterol esters were quantified by using a cholesterol assay kit (Fluorometric) from Invitrogen (Waltham, MA, USA). TAGs purified by TLC from the lipid extract of *Arabidopsis* leaves were quantified by using a colorimetric assay as described. ^62^

### Gas chromatography–mass spectrometry (GC-MS) analysis of phytosterol esters and phytosterols

Materials were obtained from 30-day-old plants. Each lipid extraction was from plant tissues with cholesteryl stearate, tripentadecanoin, methyl heptadecanote, and ergosterol as the internal standard (IS), acquired from Sigma (St. Louis, MO, USA). Individual phytosterol ester or free phytosterol samples were scraped from the TLC plate. The phytosterol ester fraction was saponified to free phytosterols by using 1N methanolic KOH, and the reaction was performed at 80°C for 2 h. After saponification, n-hexane was added to extract phytosterols, and the n-hexane phase was dried under a stream of nitrogen. For GC-MS analysis, de-esterificated phytosterols were converted into N,O-bis(trimethylsilyl)trifluoroacetamide content at 1% trimethylchlorosilane derivatives. The free phytosterol fraction was directly converted into derivatives for GC-MS analysis. The derivative trimethylsilyl (TMS)-phytosterols was used for targeted analysis by selected ion monitoring (SIM). The derivatives were injected into an Agilent Technologies 5975C gas chromatography–mass spectrometer equipped with a DB-5MS (UI) chromatographic column (30 m × 0.25 mm × 0.25 μm). The injection temperature and volume were set to 280°C and 2 μL, respectively. Helium was used as a carrier gas, with a 1:30 injector split. The gas chromatography conditions were as follows: injector temperature, 250°C; oven temperature programme, 1) 200°C for 1 min, then increasing at 10°C min^−1^ to 250°C, and maintaining this temperature for 2 min, 2) 10°C min^−1^ to 270°C and maintaining this temperature for 27 min, and 3) 10°C min^−1^ to 280°C and maintaining this temperature for 5 min.

### Chemical treatment on agar

More than 20 seedlings (wild type and *gsease*) were grown on solidified half Murashige and Skoog agar containing 10 nM squalene, 10 nM sitosterol, 10 nM campesterol, 10 nM epi-BR, or 25 µM 20ikini. DMSO is a solvent used to dissolve squalene, campesterol, epi-BR, and bikinin. Treated samples were submitted for confocal microscope analysis. Campesterol, epi-BR, and bikinin were acquired from Sigma (St. Louis, MO, USA).

### Immunoblot analysis

Seedlings at 5 dpg were frozen in liquid nitrogen and homogenised in an MAPK activity extraction buffer (100 mM HEPES [pH 7.5], 5 mM EDTA, 5 mM EGTA, 10 mM NaF, 10 mM Na3VO4, 50 mM β-glycerophosphate, 5% glycerol, 10 mM dithiothreitol, 1 mM phenylmethylsulfonyl fluoride and 1% protease inhibitor cocktail). After centrifugation at 18,000 *g* for 40 min under 4°C, we collected the supernatant and transferred it to clean tubes. Each sample (loaded with an equal amount of total protein) was electrophoresed on 12% SDS-PAGE gel. BES1 protein levels were determined using a BES1 © antibody (R3489-2, Abiocode, USA) at 1:2000 dilution. MAPK activity was determined by immunoblot analysis with a diluted anti-phospho-ERK1/2 (pThr202/Tyr204) antibody (Sigma, St. Louis, MO, USA). Western blot analysis was performed in triplicate.

### Quantification and statistical analysis

LD quantification and western blot band intensity quantification as well as green fluorescence intensity quantification of BES1 signals were measured by ImageJ. All *P*-values were calculated by two-tailed *t* tests. *P*<0.05 was considered statistically significant.

## Acknowledgments

**General**: We thank Dr. Zhi-Yong Wang (Department of Plant Biology, Carnegie Institution for Science, Stanford, CA, USA) for generously sharing transgenic seeds (BSL2ox). We also thank Ms. Mei-Jane Fang (DNA Analysis Core Laboratory, Institute of Plant and Microbial Biology, Academia Sinica), Mr. Ji-Ying Huang (Plant Cell Biology Core Laboratory, Institute of Plant and Microbial Biology, Academia Sinica) and Ms. Yu-Ching Wu (Small Molecule Metabolomics core facility, Institute of Plant and Microbial Biology, Academia Sinica) for assistance with qRT-PCR, confocal microscopy, and GC-MS experiments, respectively. We specially acknowledge Ms. Yi-Chun Chang for artwork.

**Funding:** This research was supported by the Ministry of Science and Technology grants 106-2313-B-001-008 -MY3 to Guang-Yuh Jauh. C.-M. K. H. was supported by Academia Sinica Career Development Award (AS-CDA-111-L01) and MOST research grants (111-2311-B-001-015-MY3).

**Author contributions:** C.-C. Y., C.-M. K. H., Y.-W. H., K.-C. L., J.-F. S. and G.-Y. J. conceived and designed the experiments; C.-C. Y., S.-S. C., T.-Y. C, C.-T. J. and C. K. performed the experiments; and C.-**(A)** C. Y., C.-M. K. H. and G.-Y. J. wrote the article.

**Competing interests:** All authors declare that none of the authors have competing financial or non-financial interests as defined by Science Advances.

**Data and materials availability:** We provide a full data availability statement in the manuscript. All data needed to evaluate the conclusions in the paper are present in the paper and/or the Supplementary Materials. Additional data related to this paper may be requested from the authors.

---

**Supplementary fig. 1.**
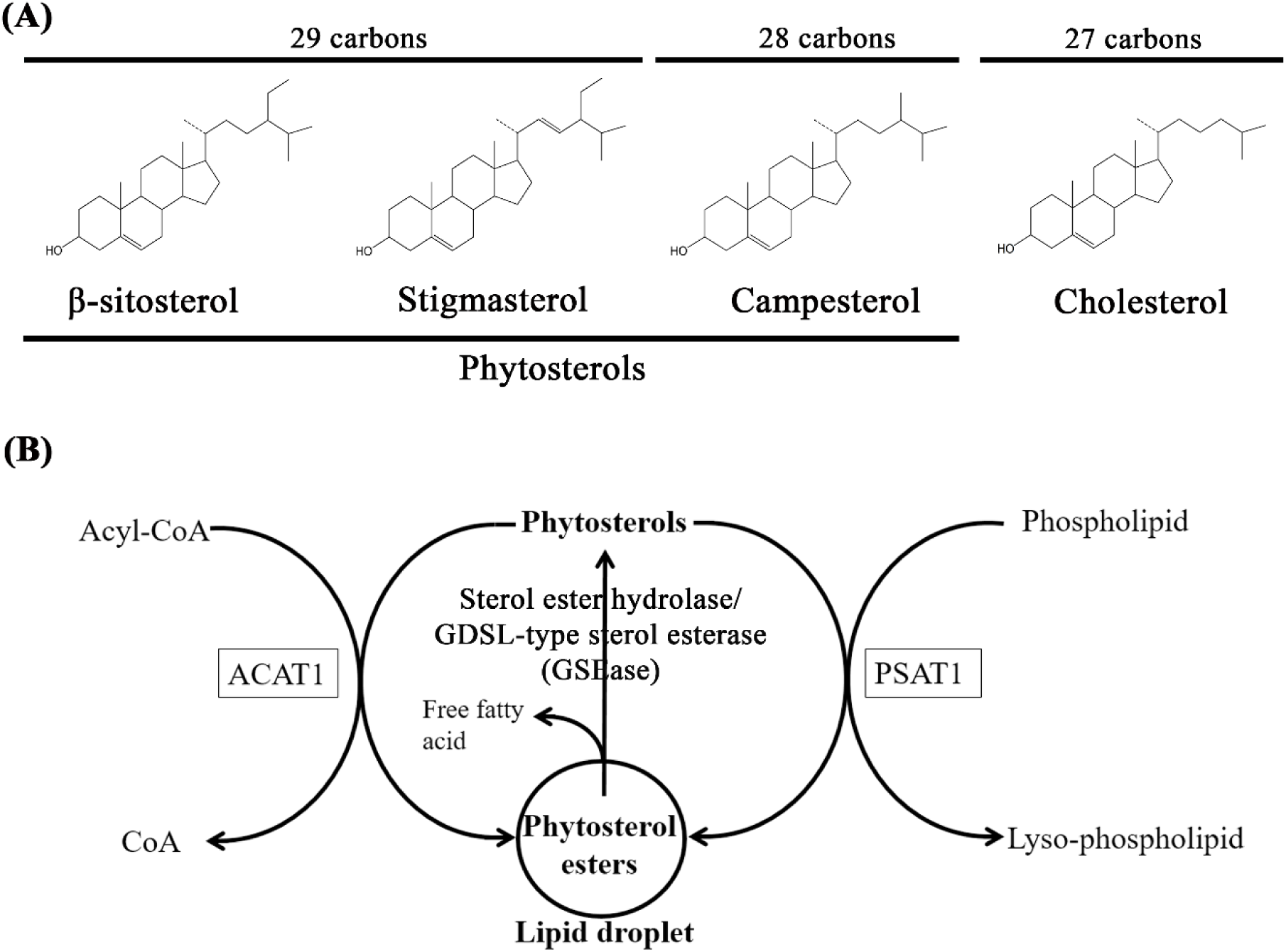
Chemical structure, distribution and homeostasis of phytosterols in plants. **(A)** Chemical structure of β-sitosterol, stigmasterol, campesterol and cholesterol. **(B)** Formation of phytosterol esters is catalysed by acyl-CoA:cholesterol acyltransferase (ACAT1) or phospholipid:sterol acyltransferase 1 (PSAT1). ACAT1 and PSAT1 catalyse phytosterol to phytosterol esters by using acyl-CoA and phospholipids, respectively, as acyl-donors ^26, 28, 29^. The phytosterol esters are stored in lipid droplets. The reaction of degrading phytosterol esters into phytosterols and a free fatty acid is *via* a sterol ester hydrolase such as GDSL-type sterol esterase (GSEase) presented in this paper.

**Supplementary fig. 2.**
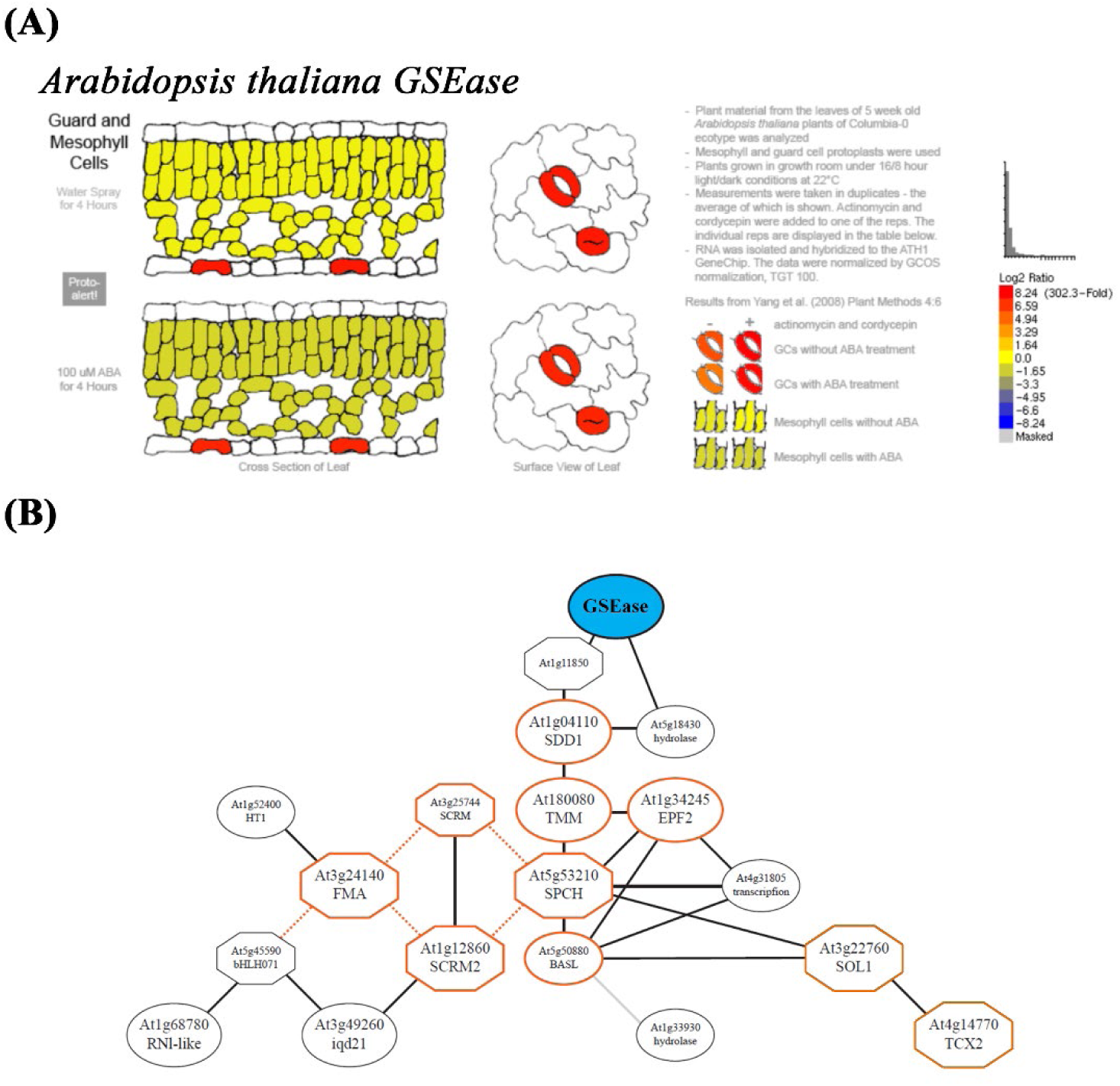
*In silico* analysis of *GSEase*. **(A)** Tissue-specific expression of *Arabidopsis thaliana GSEase* from Arabidopsis eFP Browser (http://bar.utoronto.ca/efp/cgi-bin/efpWeb.cgi). **(B)** *GSEase* coexpression network predicted by ATTED-II. *GSEase* (blue shaded oval) was used as a query in the ATTED-II database (http://atted.jp). The output data indicated that 18 genes were coexpressed with *GSEase*. Genes marked in red are stomatal development genes, such as *SDD1*, *TMM*, *EPF2*, *SPCH*, *SCRM*, *SCRM2*, *FMA*, *BASL, SUPPRESSOR OF LLP1 1* (*SOL1*) and *TESMIN/TSO1-LIKE CXC 2* (*TCX2*). Octagons indicate transcription factors, and ovals indicate genes other than transcription factors.

**Supplementary fig. 3.**
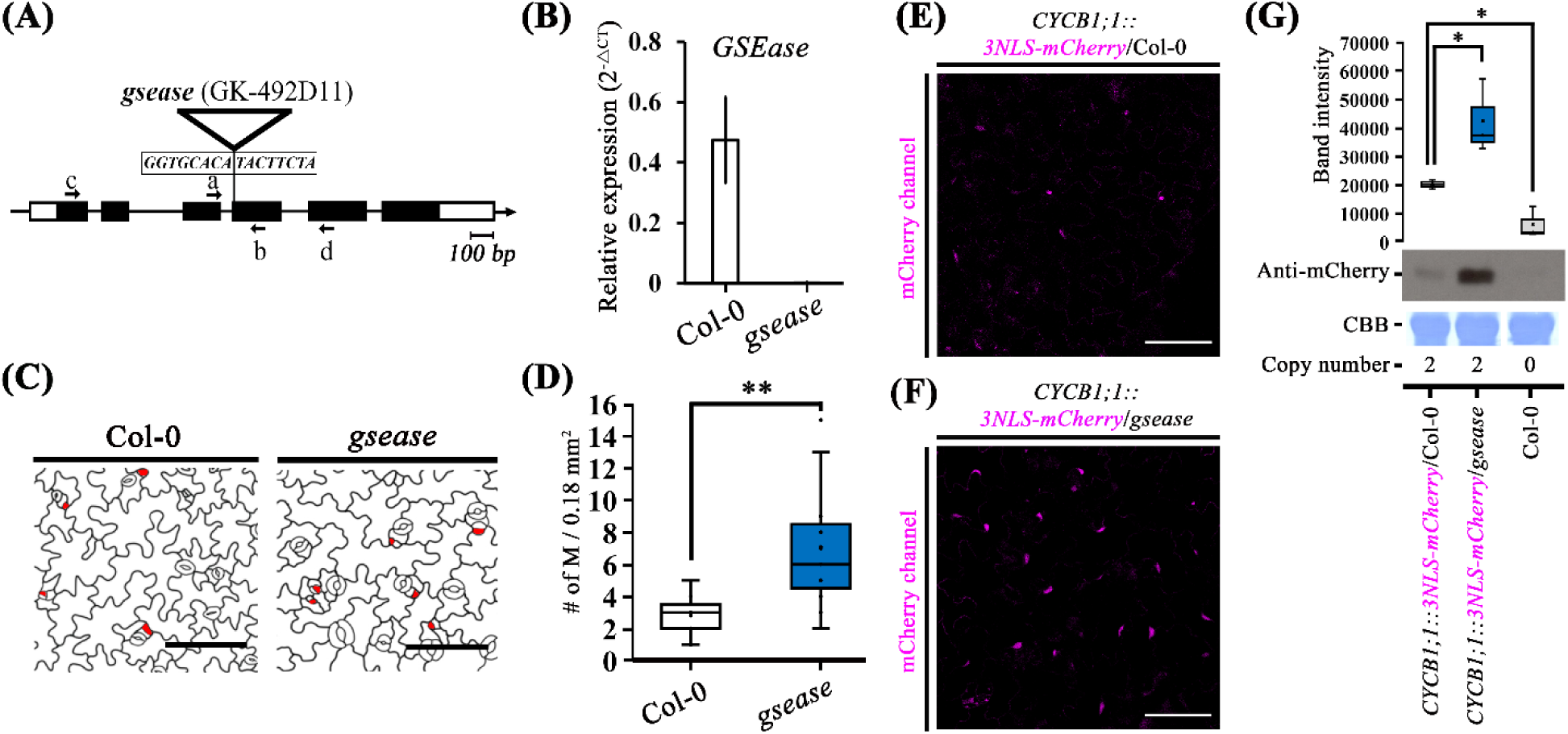
Lack of *GSEase* increases meristemoid (M) density and cell division. **(A)** Illustration of *At1g33811* locus with the exon (black bar), intron (line), and untranslated region (UTR) (white bar). T-DNA inserted site is indicated by triangle. Primers used for qRT-PCR are indicated by a and b arrows and for RT-PCR primer binding sites by c and d arrows. **(B)** qRT-PCR analysis of transcript levels of *GSEase* in wild type (Col-0) and *gsease* mutant. Data were normalised to the moderately stable reference gene *PROTEIN PHOSPHATASE 2A SUBUNIT A3* (*PP2AA3*). Data are mean ± SD of four independent experiments. **(C)** Fake images of 3-dpg abaxial cotyledon with propidium iodide staining. Ms are in red. **(D)** Quantification of Ms density in 3-dpg cotyledons in Col-0 and *gsease* leaves. The results were obtained from 11 independent lines, each independent plant involving one picture. \*\**P*<0.01. Data are median (horizontal line), range (box edges) and interquartile range (whiskers). **(E-F)** Confocal images of 5-dpg cotyledons expressing *CYCB1;1pro*::*3NLS-mCherry* in Col-0 **(E)** or *gsease* **(F)**. The 3NLS-mCherry is labelled with magenta. **(G)** Protein abundance of mCherry monitored by western blot analysis in Col-0 and *gsease* with mCherry antibody (GTX128508, GeneTex, USA). Total protein was visualised by Coomassie brilliant blue (CBB) staining. Data are median (horizontal line), range (box edges) and interquartile range (whiskers). Note, the copy number of *CYCB1;1*::*3NLS-mCherry*/Col-0 and *CYCB1;1*::*3NLS-mCherry*/*gsease* transgenic plant was equal. Copy number determination was referenced from Kanwar et al. ^63^. Data in **(B)** were normalised to the moderately stable reference gene *PP2AA3*. Scale bars are 100 µm in **(C)**, **(E)** and **(F)**. \*\**P*<0.01 and \**P*<0.05. *P*-values were calculated using one-way ANOVA with Tukey honestly significant difference test.

**Supplementary fig. 4.**
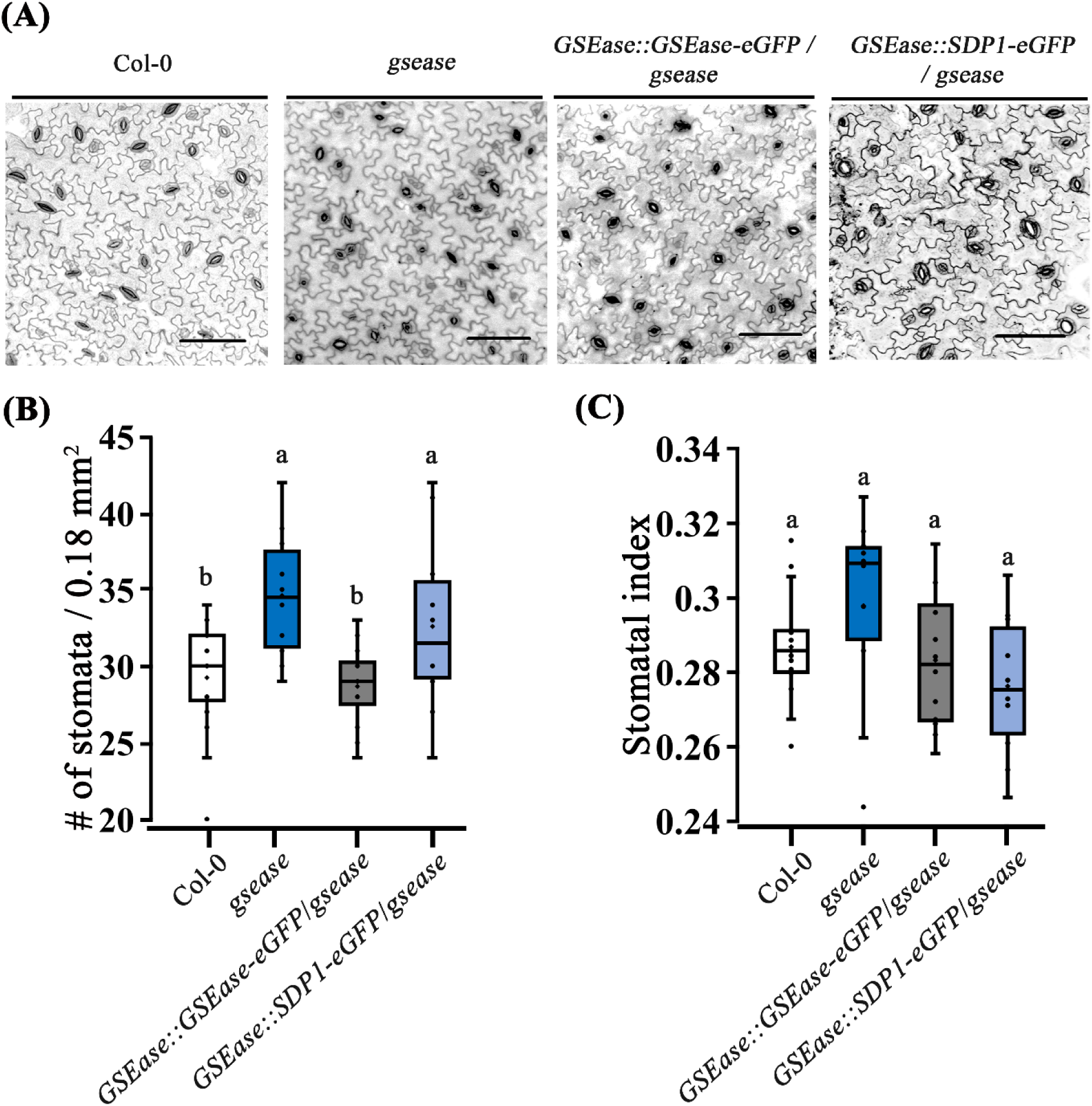
Stomata phenotype of *gsease* is complemented by expression of GSEase. **(A)** Confocal images of Col-0, *gsease* and *GSEase promoter-driven GSEase* fusing *eGFP* (*GSEase::GSEase– eGFP*) used to complement the *gsease* phenotype. *GSEase promoter-driven SUGAR-DEPENDENT1* (*SDP1*) fused with *eGFP* was a negative control. The cell outline was visualised with propidium iodide. **(B-C)** Box plots of the quantification of stomatal phenotype, stomatal density **(B)** and stomatal index **(C),** of *gsease* and its complementation lines. Data are median (horizontal line), range (box edges) and interquartile range (whiskers). The results were from 16, 10, 12 and 10 independent lines. Each independent plant described in **(B)** and **(C)** involved only one picture. Scale bars are 100 µm in **(A)**. *P*-values were calculated using one-way ANOVA with Tukey honestly significant difference test.

**Supplementary fig. 5.**
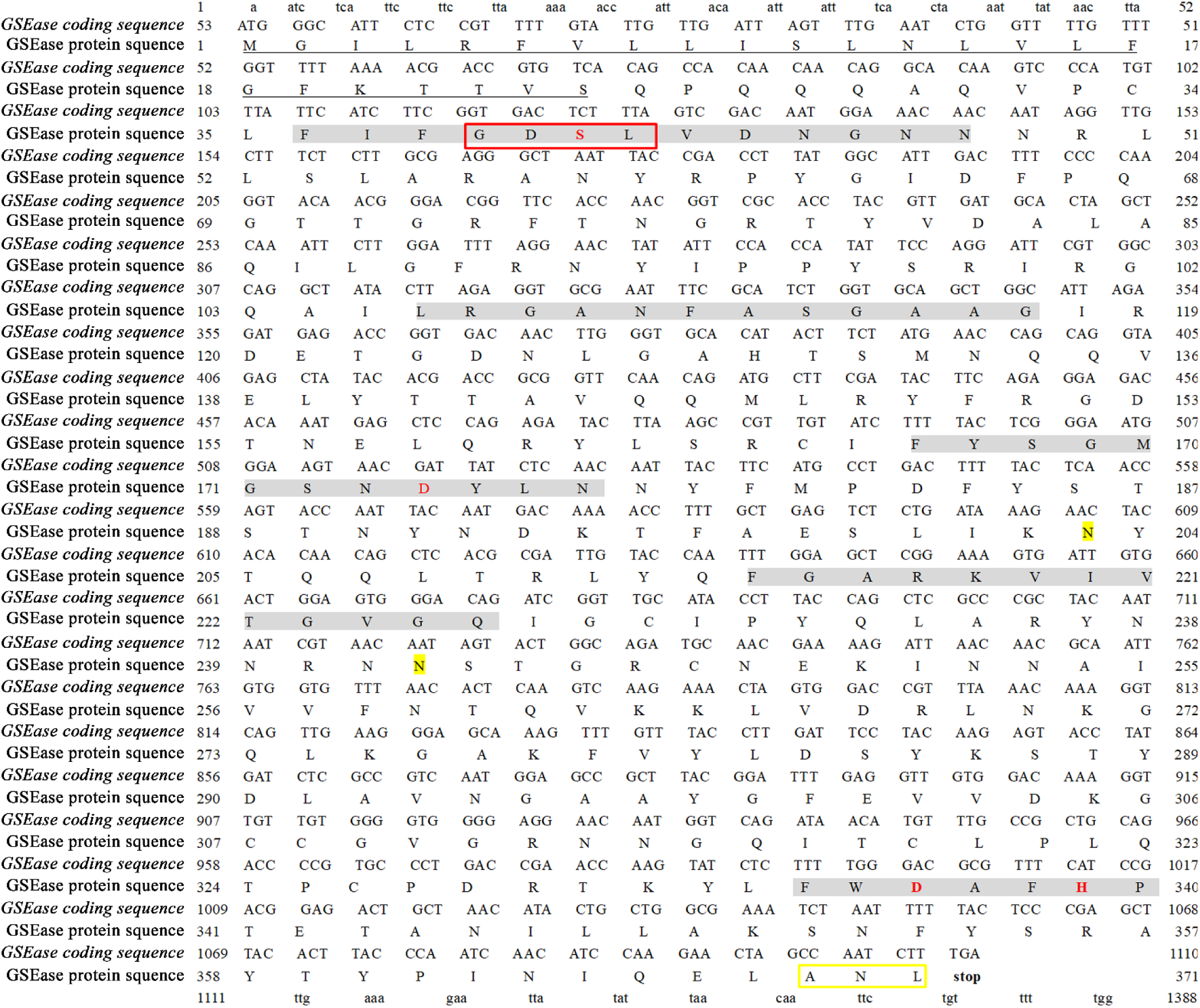
GSEase coding sequence and protein sequence. The GDSL family has five conserved blocks (Ⅰ–Ⅴ, highlighted in grey) ^66^. The conserved motif locates in the first block (labelled in a red box). The putative active site residues (S41, D174, D336, and H339) are marked in red. The underlined sequence was predicted to be a signal peptide (SP) by the SignalP 5.0 server (http://www.cbs.dtu.dk/services/SignalP/). Putative N-glycosylation sides (N203 and N242) are labelled in yellow according to the NetNGlyc 1.0 Server (http://www.cbs.dtu.dk/services/NetNGlyc/). The putative peroxisome targeting signal type 1 tripeptide is enclosed in the yellow box ^67^.

**Supplementary fig. 6.**
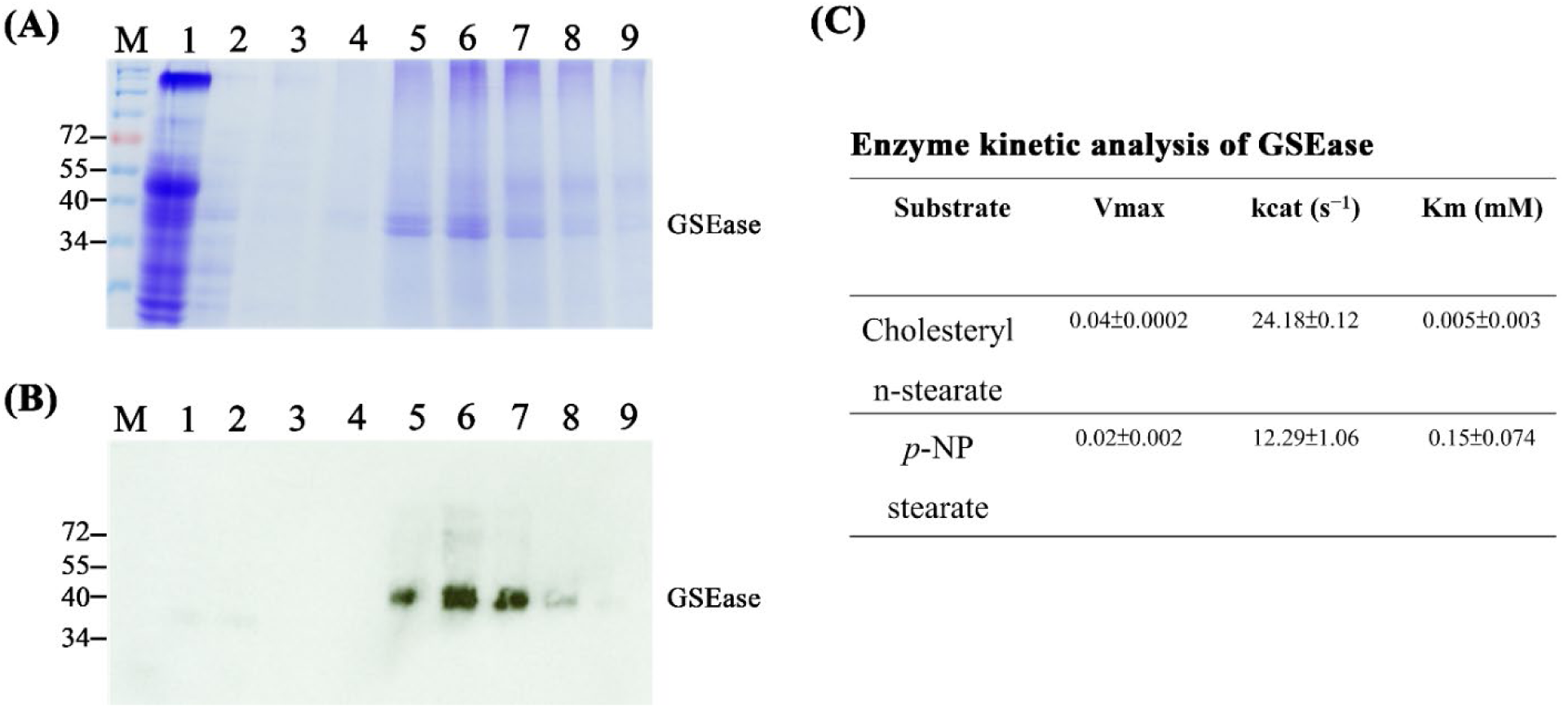
Purification of histidine-tagged GESase (GSEase-6xHis) and detection of *GESase* kinetic analysis. Overexpressed GSEase in an *Escherichia coli* or in a *Pichia pastoris* system was not effective in our experimental setup. Therefore, we generated a GSEase CDS (**SFig. 3**) overexpression line using the *Pro35S::GSEase CDS-6x histidine* in *A. thaliana*. GSEase recombinant proteins were then purified with diethylaminoethanol (DEAE) sepharose and the histidine-tag-column procedure and verified by western blot assay. The purified protein was then used for characterising its substrate specificity *in vitro*. **(A-B)** Purification of histidine-tagged GESase (GSEase-6xHis). SDS-PAGE **(A)** and western blot **(B)** detection of the GSEase-6xHis during purification steps. M, Precision Plus Protein Standards Dual Color, BIO-RAD; lane 1, crude protein extract collected from DEAE column elution; lane 2, flow through; lane 3, 20 mM imidazole; lane 4, 40 mM imidazole; lane 5, 60 mM imidazole; lane 6, 80 mM imidazole; lane 7, 100 mM imidazole; lane 8, 120 imidazole; lane 9, 140 mM imidazole. Western blot analysis involved using a monoclonal 6X His tag antibody (GT359, GeneTex, USA). **(C)** Enzyme kinetic analysis of GSEase**-**6xHis indicating that GSEase processed a higher (≈2 times) maximum rate of production (Vmax) and turnover number (kcat) in cholesteryl n-stearates than in *p*-NP stearate. Furthermore, the higher substrate affinity of cholesteryl n-stearate is indicated by its lower Km value than that of *p*-NP stearate. Lipids possess an acyl and alcohol group, and both fit in their respective binding tunnel in a lipase ^64^. *p*-NP stearate and cholesteryl n-stearate have the same acyl group but different alcohol groups. Thus, the substrate affinity of GSEase is possibly determined on the alcohol group binding tunnel but not the acyl group binding tunnel. In addition, we assumed that the alcohol group binding tunnel explains why GSEase does not prefer triacylglycerol (TAG) substrates such as tristearate because the alcohol group of TAGs may not be suitable for GSEase binding. The kinetic parameters were calculated with the Lineweaver–Burk equation in a nonlinear regression, with cholesteryl n-stearate or *p*-NP stearate used as substrates at room temperature at pH 8.0. Data are mean ± SD of three independent enzyme reactions.

**Supplementary fig. 7.**
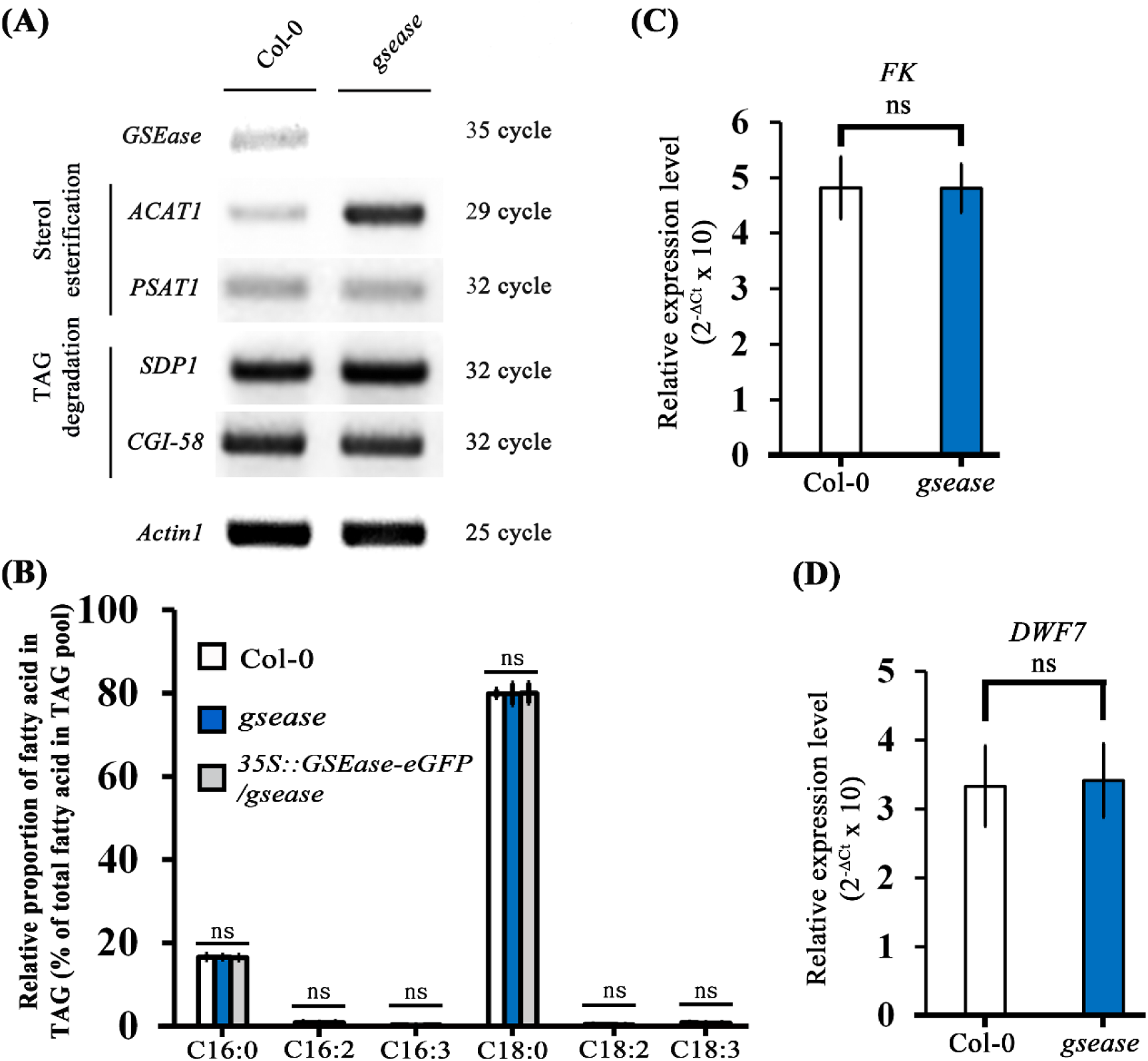
Lack of *GSEase* does not affect triacylglycerol (TAG) metabolism or phytosterol synthesis. **(A)** RT-PCR analysis of transcript levels of sterol esterification genes (*acyl-CoA:cholesterol acyltransferase 1* [*ACAT1*] and *phospholipid:sterol acyltransferase 1* [*PSAT1*]) and TAG-degradation genes, *SDP1* and *comparative gene identification-58* (*CGI-58*), in Col-0 and *gsease*. The moderately stable reference gene *Actin1* was a control. **(B)** Fatty acid composition of TAGs in Arabidopsis leaves of Col-0, *gsease* and 35S::GSEase:eGFP/*gsease* by GC-MS. In briefly, TAG samples were collected from TLC separation. Samples were added to 1 ml 2.5% H_2_SO_4_ (v/v) in MeOH and incubated at 80°C for 1 h to form fatty acid methyl esters (FAMEs). After cooling to room temperature, sample extracts were pooled and added to 500 µl n-hexane and 1.5 ml 0.9% NaCl (w/v) to separate two phases. The upper phase containing FAMEs underwent GC-MS analysis with Agilent Technologies 5975C GC/MS equipped with a DB-5MS (UI) chromatographic column (40 m x 250 μm x 0.25 μm). The split ratio was 1:30 and the injector temperature 250 °C. The column temperature program was initially held at 150 °C for 3 min; ramped from 150 to 240 °C at for 9 min; and held at 240 °C for 5 min. The standard F.A.M.E. Mix C8-C24 was acquired from Sigma (St. Louis, MO, USA). Tripentadecanoin was used as the internal standard (IS). Data are mean ± SD of three independent experiments. NS = not significant. **(C-D)** Phytoterol biosynthesis genes levels were not altered in *gsease*. qRT-PCR analysis of transcript levels of *FK* **(C)** and *DWF7* **(D)** in Col-0 and *gsease*. Data in **(C)** and **(D)** were normalized to the moderately stable reference gene *PP2AA3*. Data are mean ± SD of three independent experiments. NS = not significant. *P*-values were calculated using one-way ANOVA with Tukey honestly significant difference test.

**Supplementary fig. 8.**
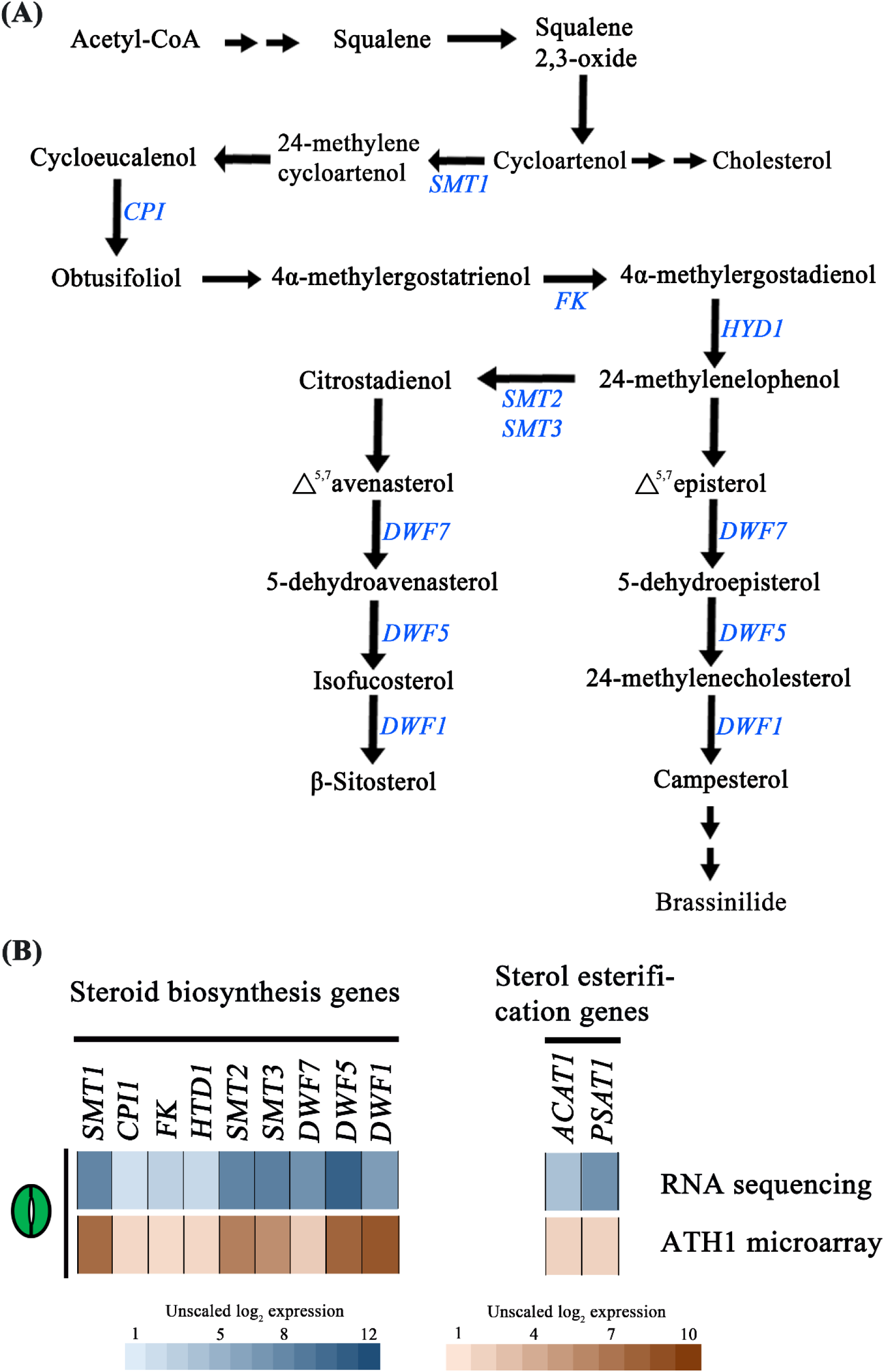
Metabolic chart of steroid biosynthesis and expression of corresponding genes in mature guard cells (GCs). To evaluate whether the steroid biosynthesis and sterol esterification pathway are present in GCs, we checked some key enzymes related to the steroid biosynthesis and sterol esterification pathway according to transcriptome data ^31^. **(A)** Simplified steroid biosynthesis pathway. The steroid biosynthesis genes marked in blue are *STEROL METHYLTRANSFERASE 1* (*SMT1*), *CYCLOPROPYL ISOMERASE* (*CPI1*), *STEROL 14a-DEMETHYLASE/CYTOCHROME P450 51A2* (*CYP51A2*), *FACKEL* (*FK*), *HYDRA1* (*HYD1*), *STEROL METHYLTRANSFERASE 2* (*SMT2*), *STEROL METHYLTRANSFERASE 3* (*SMT3*), *DWARF 7 (DWF7)*, *DWARF 5 (DWF5)*, and *DWARF 1(DWF1)*. **(B)** Expression of the steroid biosynthesis and sterol esterification pathway genes in mature GCs obtained by RNA sequencing and an ATH1 microarray in Adrian et al. ^31^. All selected genes were related to phytosterol synthesis and esterification expressed in GCs.

**Supplementary fig. 9.**
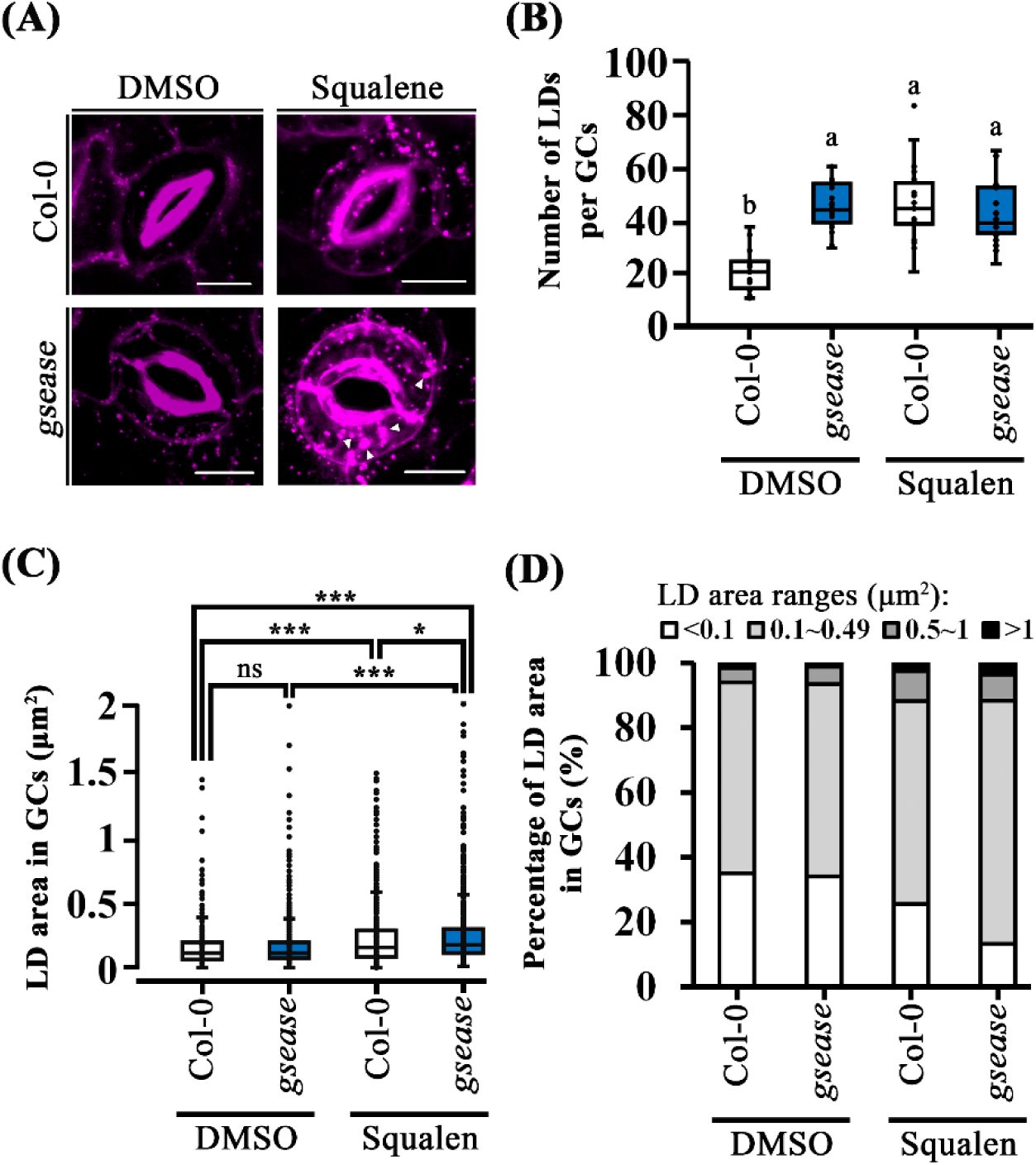
Phytosterol precursor, squalene, treatment barely increased the number but enlarged the size of lipid droplets (LDs) in stoma. **(A)** Confocal images of LDs of GCs in Col-0 and *gsease* with or without squalene treatment. LDs were stained with nile red. The LD number was significantly increased in the GCs of Col-0 grown on a solidified half Murashige and Skoog agar containing 10 nM squalene. White triangle indicated enlarged LD in GCs of *gsease* after squalene treatment. **(B-C)** LD number **(B)** and LD area **(C)** in GCs from Col-0 and *gsease* with or without squalene treatment. LD number was greater in *gsease* than Col-0, but LD area did not differ between Col-0 and *gseas*e. After treatment with squalene, LD number in the GCs of Col-0 and *gsease* was not significantly increased, but the LD area was enlarged. **(D)** Percentage of LD area of GCs in Col-0 and *gsease* with or without squalene treatment. Percentage of LD area in GCs in Col-0 and *gsease* was similar, but squalene-treated *gsease* showed increased percentage size range about 0.1–0.49, 0.5–1 and >1 in GCs of Col-0 and *gsease*. Scale bars are 10 µm in **(A)**. The results were obtained for more than 18 cells (n>18) from 10 independent lines. *P*-values were calculated using one-way ANOVA with Tukey honestly significant difference test.

**Supplementary fig. 10.**
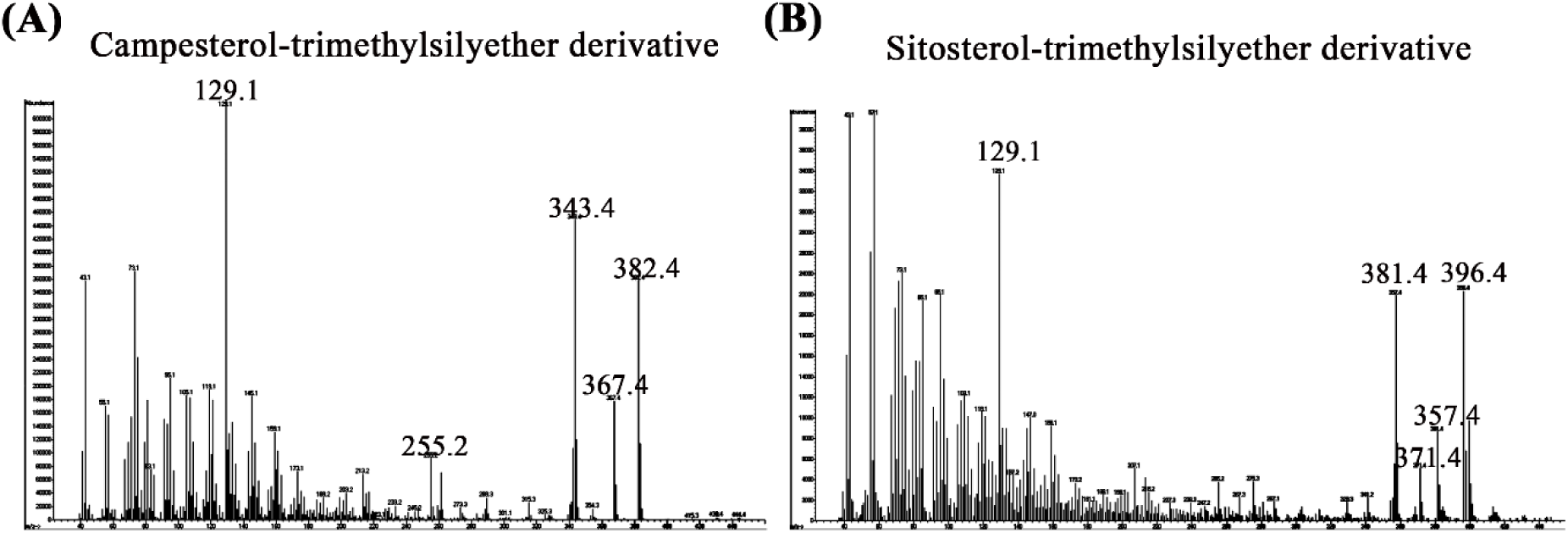
GC-MS results of campesterol- and sitosterol-trimethylsilyether derivative. Campesterol-trimethylsilyether derivative and **(A)** sitosterol-trimethylsilyether derivative **(B)**. Phytosterol derivatization methods are described in Methods.

**Supplementary fig. 11.**
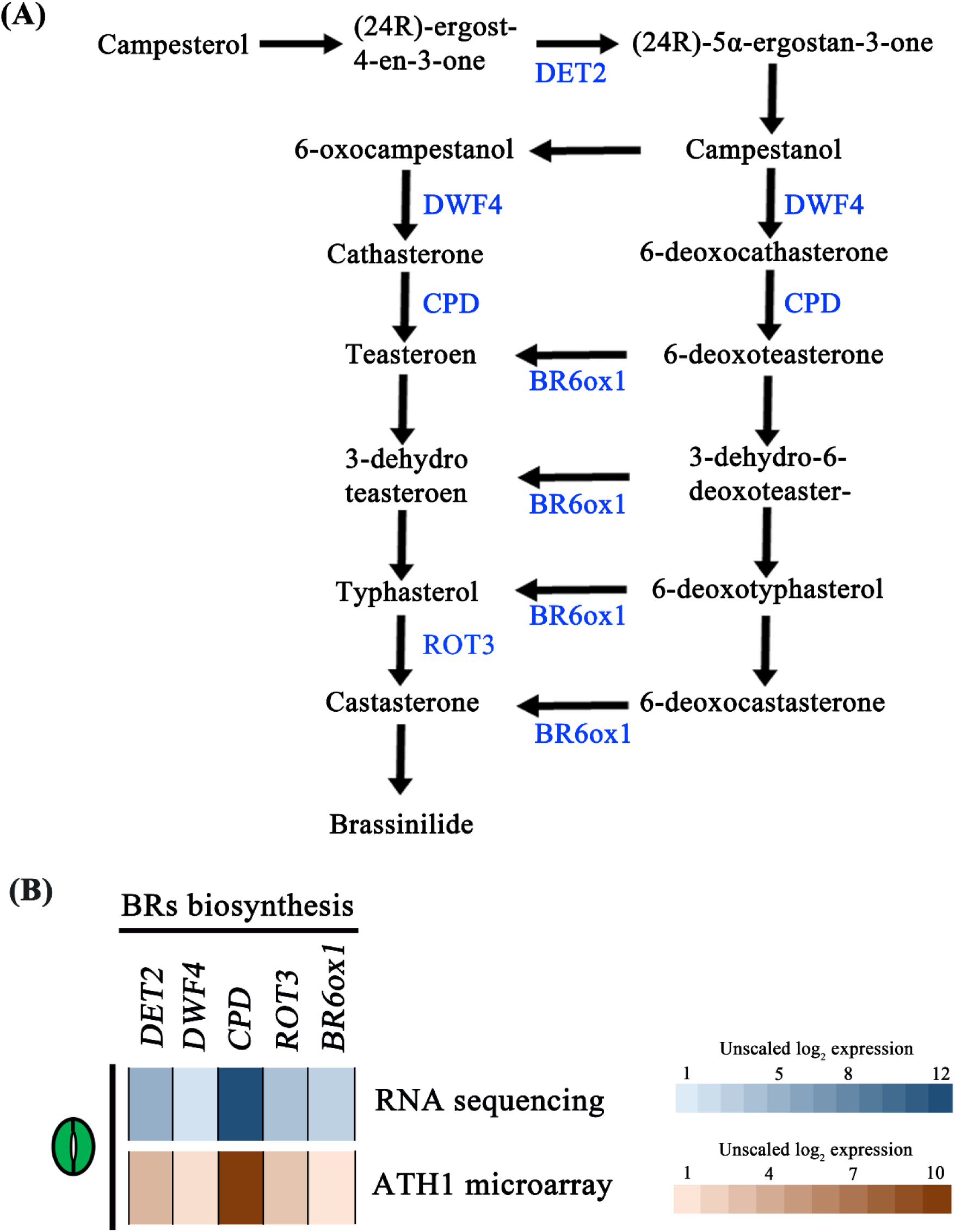
Metabolic chart and expression abundance of brassinosteroid (BR) biosynthetic genes in mature GCs. (A) Conventional proposed BR biosynthetic pathway. BR biosynthetic genes listed in blue are *DWF5*, *DWF4*, *de-etiolated2* (*DET2*), *CPD*, *BR-6-Oxidase1* (*BR6ox1*), and *rotundifolia 3* (*ROT3*). (B) Expression of the BR biosynthesis genes from mature GCs in public datasets of RNA sequencing and an ATH1 microarray ^31^.

**Supplementary fig. 12.**
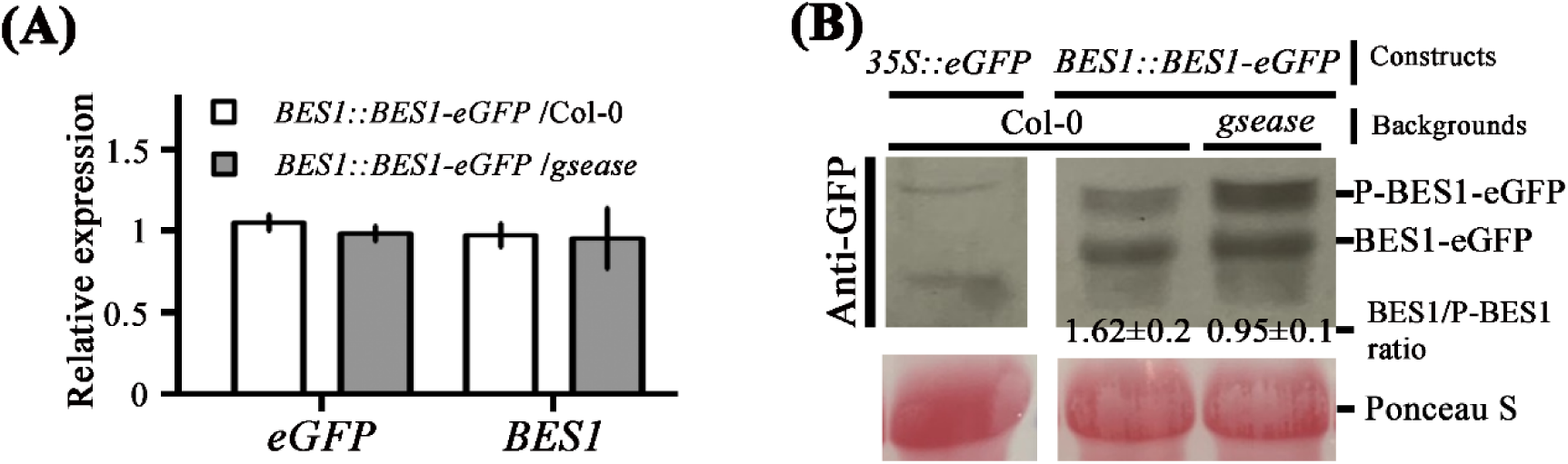
Lack of *GSEase* reduced brassinosteroid (BR) response post-translationally. **(A)** Transcriptional levels of *eGFP* and *BRI1-EMS SUPPRESSOR1 (BES1)* genes in seedlings at 4 days post-growth (dpg) seedlings from BES1::BES1-GFP/Col-0 and BES1::BES1-GFP/*gsease*. Data were normalized to the moderately stable reference gene *Actin1*. **(B)** Western blot analysis of BES1-GFP in 4-dpg Col-0 and *gsease* seedlings using GFP antibody (GTX113617, GeneTex, USA). Total protein was visualized with Ponceau S staining. The numbers are the ratio of levels of unphosphorylated BES1 (BES1) to phosphorylated BES1 (P-BES1). Data are mean ± SD of three independent experiments. *P*-values were calculated using one-way ANOVA with Tukey honestly significant difference test.

**Supplementary fig. 13.**
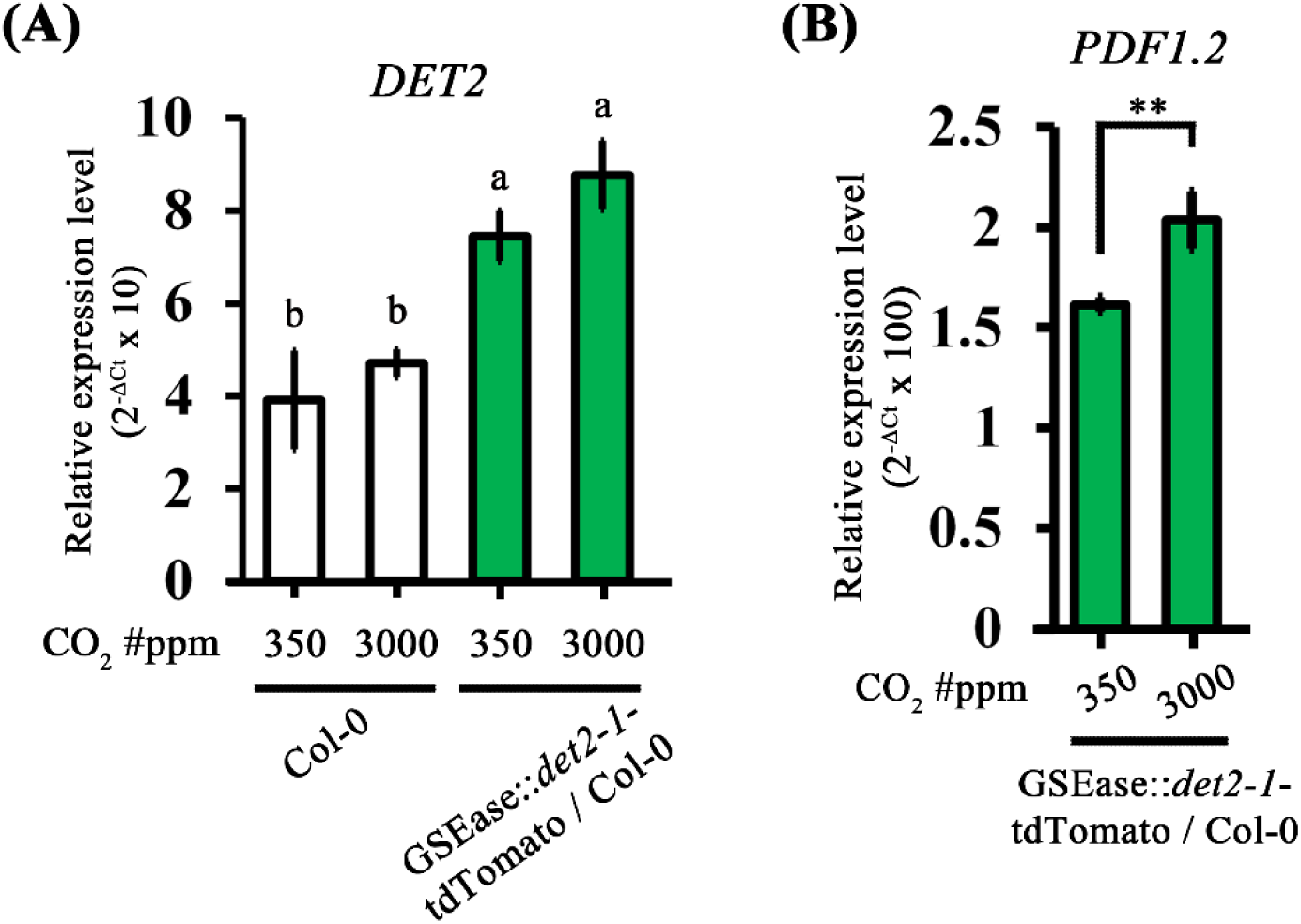
Transcript levels of *DET2* and *PDF1.2* (a CO_2_ inducible gene) in Col-0 and GSEase::*det2-1*-tdTomato / Col-0 under normal (350 ppm) and high CO_2_ condition (3000ppm). **(A)** Transcript levels of *DET2* in Col-0 and GSEase::*det2-1*-tdTomato / Col-0 were not affected under high CO_2_ condition versus the normal condition. **(B)** Transcript levels of *PDF1.2*, a CO_2_ inducible gene ^65^, in GSEase::*det2-1*-tdTomato / Col-0 were upregulated under high CO_2_ versus the normal condition. Data are mean ± SD from three independent experiments. *P*-values were calculated using one-way ANOVA with Tukey honestly significant difference test.

**Supplementary fig. 14.**
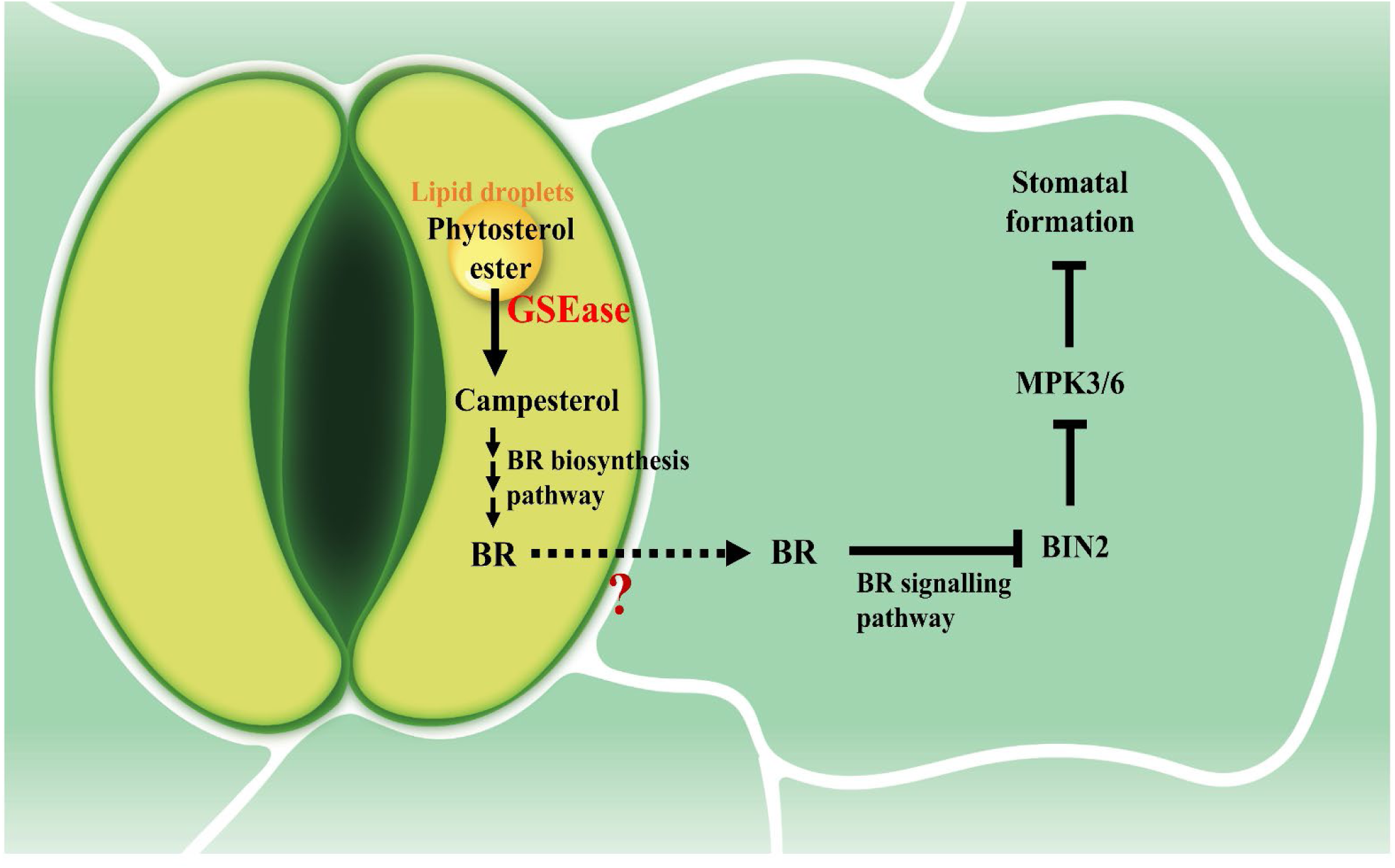
A hypothetical model of GC phytosterol homeostasis in regulating stomata formation. GSEase, a stomatal specific sterol esterase, plays a critical role in hydrolysing storage-formed phytosterol esters from LDs in GCs. The liberated phytosterols (campesterol) function as a precursor of BRs, mediating its downstream BR signaling to prevent stomata formation in adjacent cells. Dotted lines reveal the unclear question: how BRs are transported from GCs to adjacent cells.

**Supplementary tab. 1.** Neutral lipid content in leaves of Col-0, *gsease*, and 35S::GSEase-GFP/*gsease* plants.

| Genotypes | Squalene | Phytosterol esters<br>(mg/g dry weight) | Free phytosterols<br>(mg/g dry weight) | TAGs (mg/g dry<br>weight) |
| --- | --- | --- | --- | --- |
| Col-0 | - | 0.39±0.06 (100%<sup>a</sup>) | 2.15±0.12 (100%) | 0.89±0.01 (100%) |
| *gsease* | - | 0.86±0.08 (221%) | 1.6±0.20 (74%) | 0.87±0.02 (98%) |
| 35S::GSEase-GFP/*gsease* | - | 0.31±0.03 (80%) | 1.95±0.07 (91%) | 0.89±0.01 (100%) |
| Col-0 | + | 2.56±0.28 (100%) | 4.33±0.16 (100%) | 0.91±0.03 (100%) |
| *gsease* | + | 3.82±0.25 (149%) | 3.43±0.03 (79%) | 0.91±0.04 (100%) |
| 35S::GSEase-GFP/*gsease* | + | 2.6±0.28 (102%) | 4.27±0.41 (99%) | 0.89±0.04 (98%) |

<sup>a</sup>The relative percentage of each sample is in parentheses. Col-0 values are defined as 100%.
Data are mean±SD of three independent experiments. TAGs, triacylglycerols

**Supplementary tab. 2.** Phytosterol profiles in Col-0 and *gsease* rosette leaves.

| Form | Phytosterols | Col-0 (µg/g dry weight) | *gsease* (µg/g dry weight) |
| --- | --- | --- | --- |
| Free form | Free sitosterol | 1816.76±97.7 (100%<sup>a</sup>) | 1218.92±182.34 (67.1%) |
|  | Free campesterol | 104.5±27.84 (100%) | 51.8±7.58 (49.6%) |
| Storage form | Esterified sitosterol | 148.76±23.87 (100%) | 193.39±8.46 (130.0%) |
|  | Esterified campesterol | 8.86±2.78 (100%) | 17.55±1.90 (198.0%) |

<sup>a</sup>The relative percentage of each sample is in parentheses. Col-0 values are defined as 100%.
Data are mean ± SD of three independent experiments.

**Supplementary tab. 3.** Top 15 co-expressed genes of GSEase.

|  | **Genes** | **Functions** | **Other ID** |
| --- | --- | --- | --- |
| **1** | *AT1G04800* | glycine-rich protein | AT1G04800 |
| **2** | *AT5G18430* | GDSL-like Lipase/Acylhydrolase<br>superfamily protein | AT5G18430 |
| **3** | *sks9* | SKU5 similar 9 | AT4G38420 |
| **4** | *AT1G22690* | Gibberellin-regulated family protein | AT1G22690 |
| **5** | *AT4G14480* | Protein kinase superfamily protein | AT4G14480 |
| **6** | *SDD1* | Subtilase family protein | AT1G04110 |
| **7** | *AT3G62820* | Plant invertase/pectin methylesterase<br>inhibitor superfamily protein | AT3G62820 |
| **8** | *AT1G11850* | transmembrane protein | AT1G11850 |
| **9** | *ALMT12* | aluminum-activated: malate transporter 12 | AT4G17970 |
| **10** | *FMA* | basic helix-loop-helix (bHLH) DNA-binding superfamily protein | AT3G24140 |
| **11** | *HT1* | Protein kinase superfamily protein | AT1G62400 |
| **12** | *MPK12* | mitogen-activated protein kinase 12 | AT2G46070 |
| **13** | *MYB60* | myb domain protein 60 | AT1G08810 |
| **14** | *AT5G25840* | DUF1677 family protein (DUF1677) | AT5G25840 |
| **15** | *AT3G51760* | hypothetical protein (DUF688) | AT3G51760 |

Results acquired from https://atted.jp/locus/?gene_id=AT1G33811.

**Supplementary Tab. 4.**
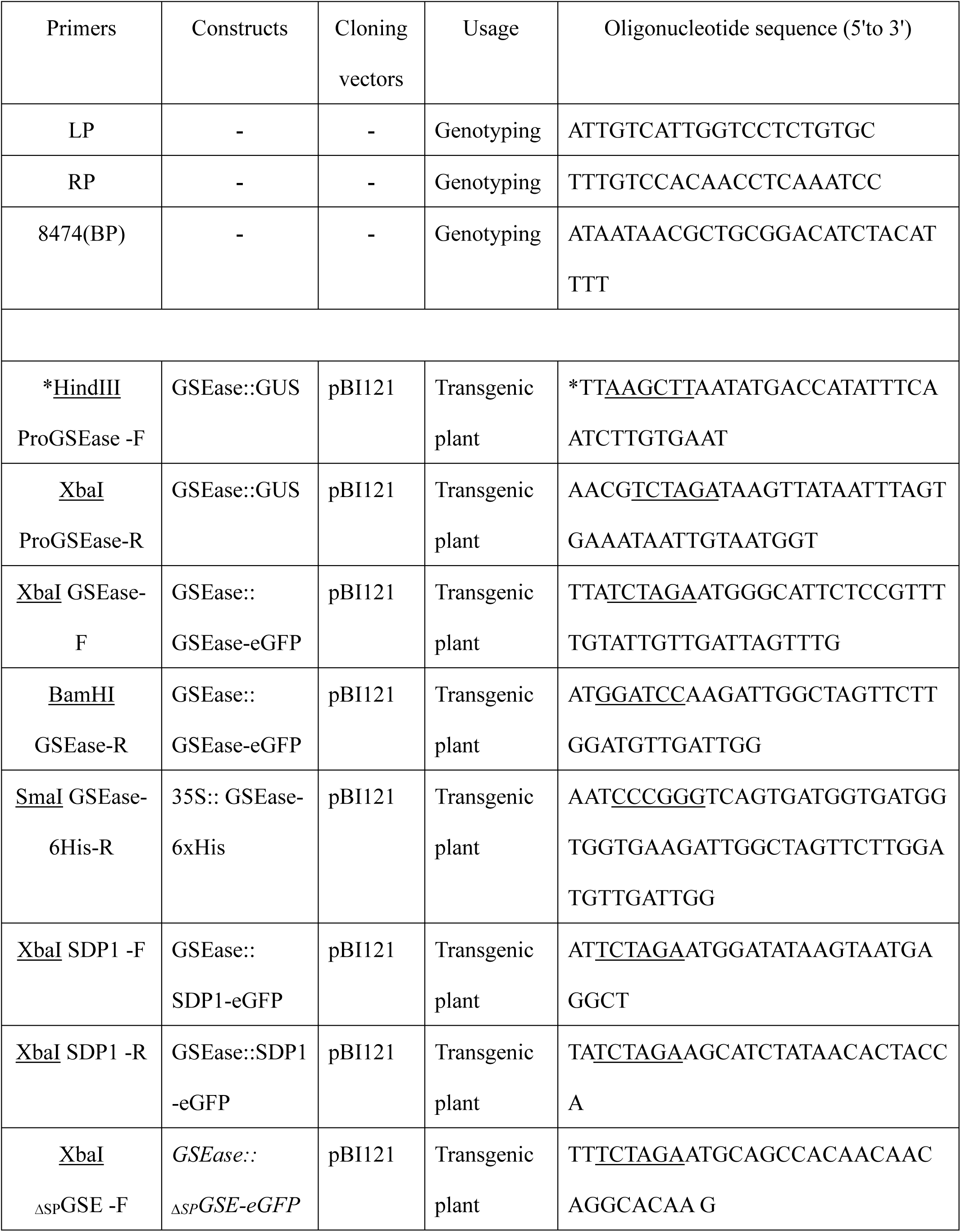
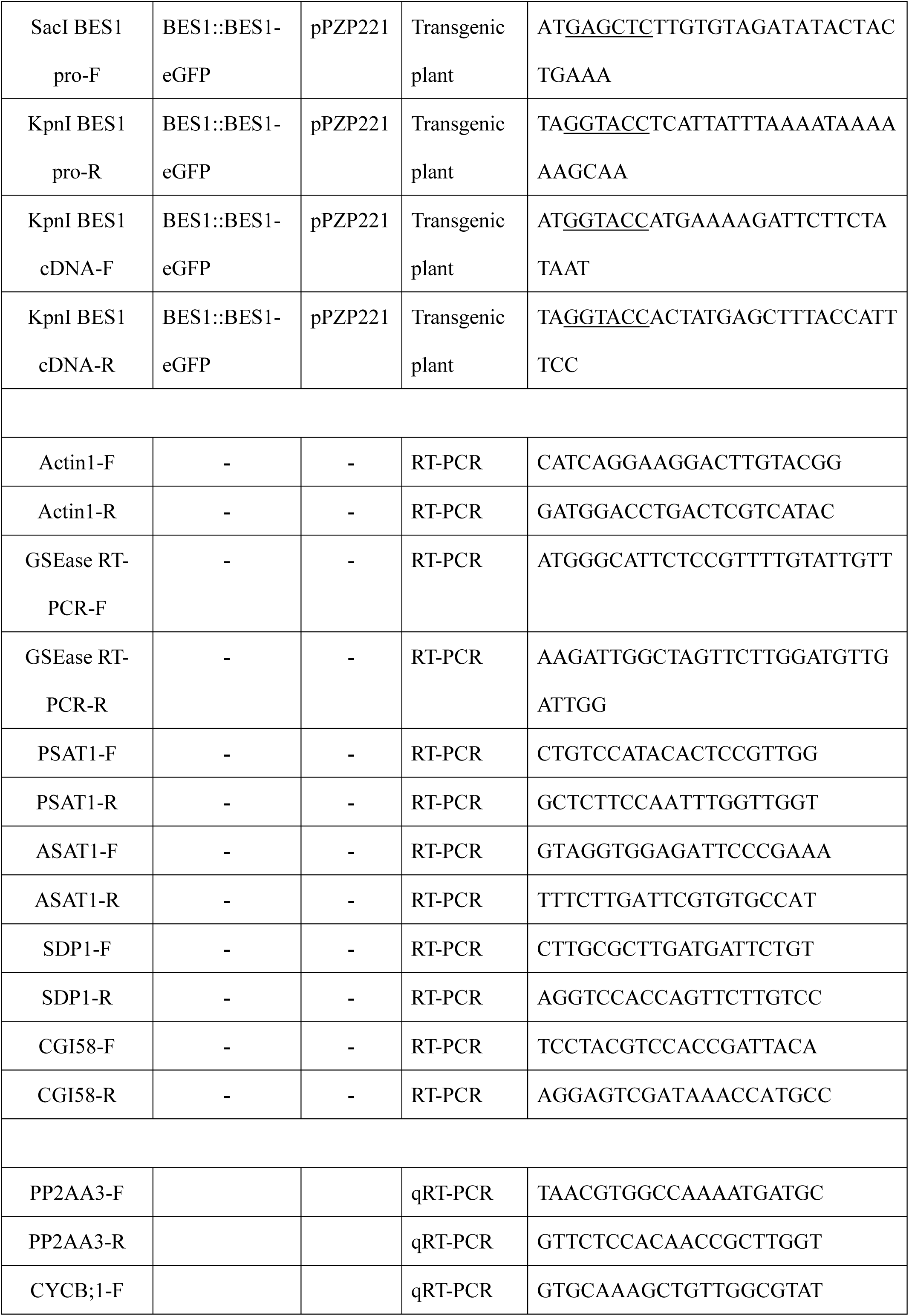
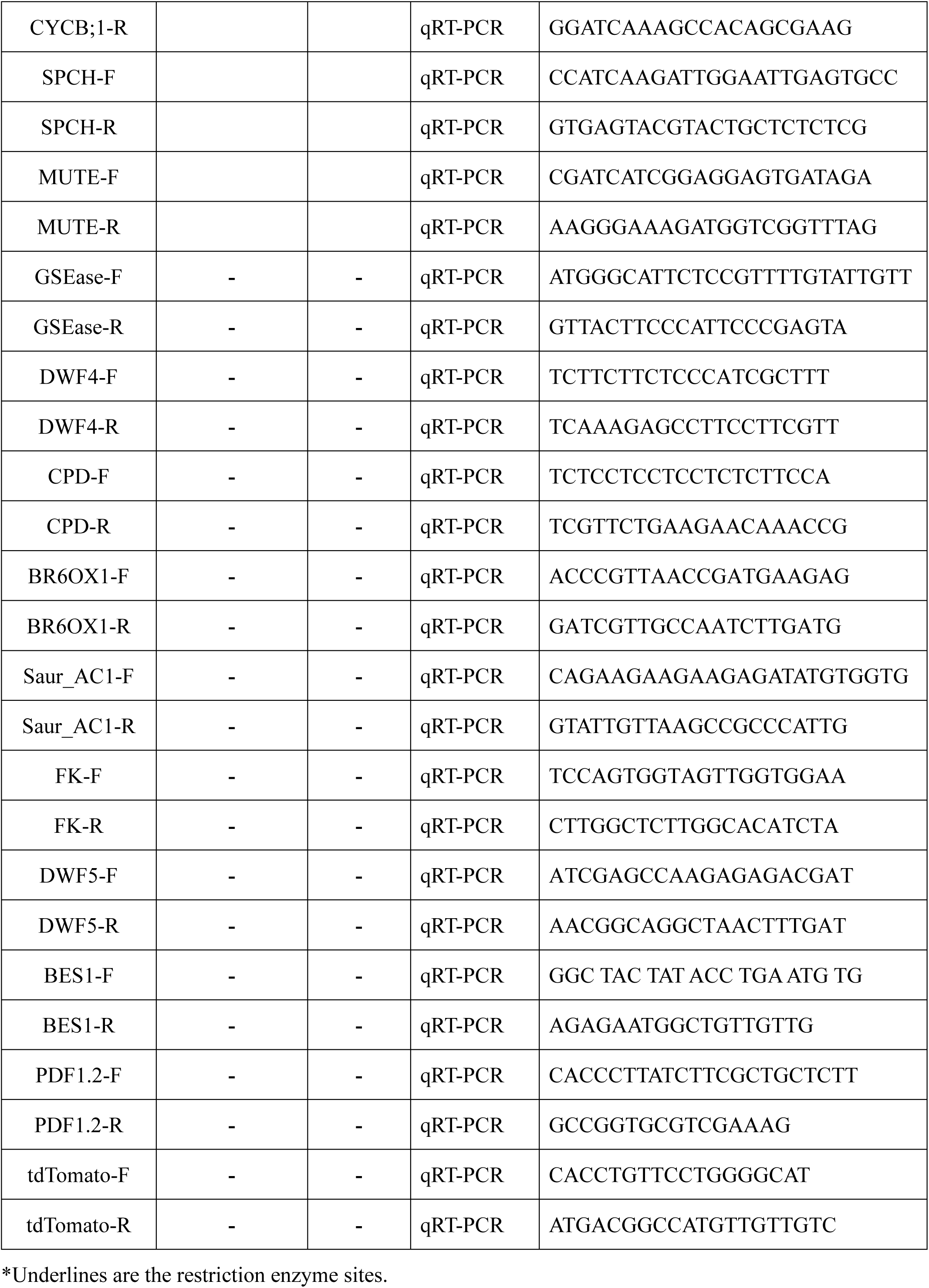
List of primers used for genotyping, GSEase localization, sub-localization assay, protein overexpression, RT-PCR and qRT-PCR.

